# Synthetic cargo adaptors reveal molecular features that enhance dynein activation

**DOI:** 10.1101/2025.06.06.658359

**Authors:** Aravintha Siva, John P. Gillies, Ashley de Borchgrave, Sharon R. Garrott, Rishi Mishra, Rita El Jbeily, Reagan S. Clarke, Cameron Conklin, Daytan Gibson, Juliana L. Zang, Morgan E. DeSantis

## Abstract

Cytoplasmic dynein-1 (dynein) facilitates the microtubule-based retrograde trafficking of all cellular cargo. To become active, dynein binds dynactin and one of many cargo-specific adaptors to form the active transport complex. Despite having similar structures, active transport complexes assembled with different adaptors move with different properties *in vitro*. To explore how adaptors differentially activate dynein, we engineered a library of synthetic adaptors and characterized their ability to activate dynein using *in vitro* reconstitution and cell-based trafficking assays. We found that the apparent motility of dynein is highly plastic and tunable by the adaptor sequence and that it is possible to engineer adaptors that outperform endogenous adaptors’ ability to generate highly motile active transport complexes. We also found that different adaptors support distinct trafficking behavior and cargo movement in cells. These findings provide insight into how dynein motility is modulated to meet the unique trafficking requirements of all cellular cargo.

## Introduction

Cytoplasmic dynein 1 (dynein) is the primary minus-end directed microtubule-based motor protein and is responsible for nearly all long-distance retrograde trafficking in most eukaryotes. Besides traditional cargo transport, dynein performs critical functions during cell division, including focusing and aligning the mitotic spindle (Raaijmakers and Medema, 2014). Mutations in dynein and proteins that regulate its activity lead to a plethora of developmental and neurodegenerative diseases (Becker et al., 2020; Marzo et al., 2019; Möller et al., 2025; Lipka et al., 2013).

Dynein is a multi-subunit protein complex assembled from dimers of six different subunits that include the heavy chain, intermediate chain, light intermediate chain and three different types of light chains (Reck-Peterson et al., 2018; Canty et al., 2021; Schmidt and Carter, 2016). The heavy chain contains a tail, an ATPase motor domain, a mobile linker domain that connects the tail and the motor domain, and a microtubule binding domain connected to the motor domain via a long coiled-coil stalk. The intermediate and light intermediate chains dock on the heavy chain tail, and the light chains are docked on a flexible region of the intermediate chain (Figure 1A) (Zhang et al., 2017).

**Figure 1:**
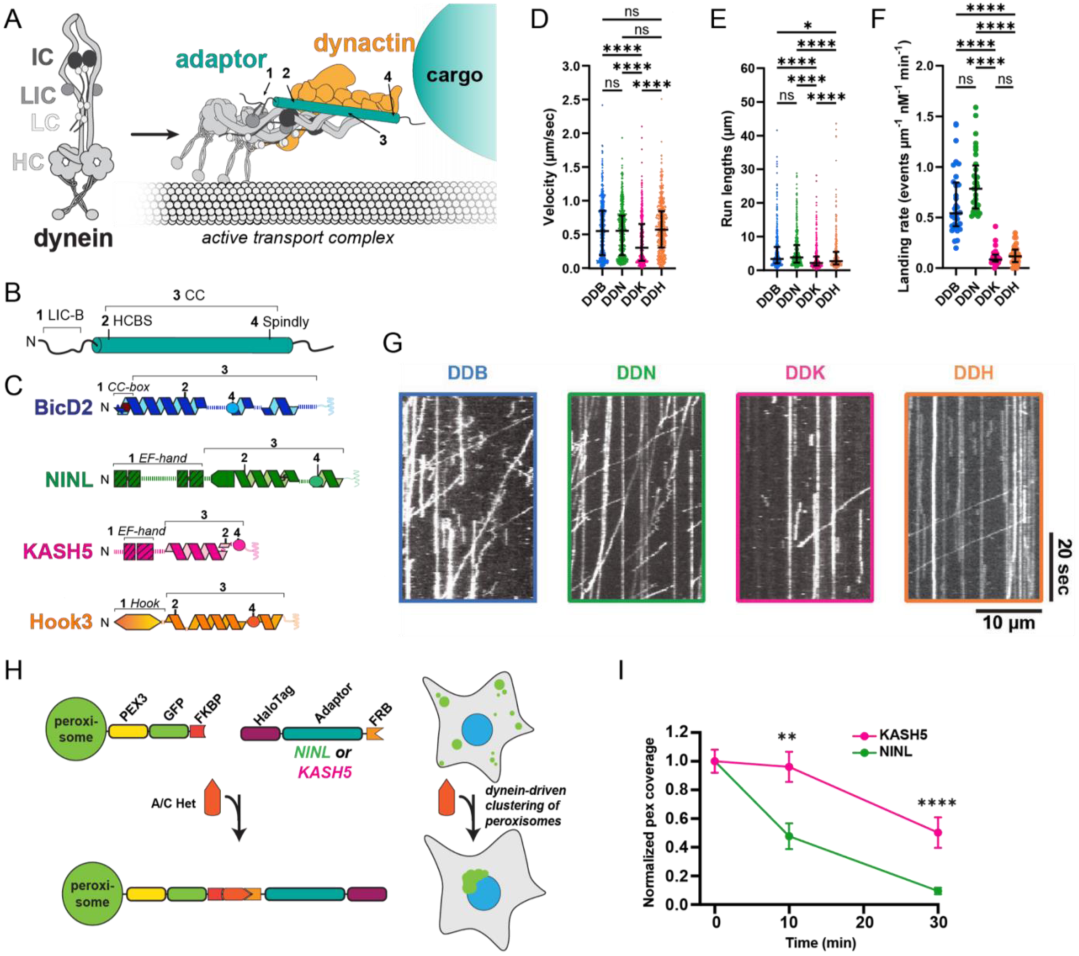
Adaptors generate active transport complexes with distinct motility. **A.** Diagram of dynein (heavy chain: HC, intermediate chain: IC, light intermediate chain: LIC and light chains: LC in different shades of gray), dynactin (yellow) and an adaptor (teal) forming an active transport complex. **B.** Schematic of a generalized adaptor with the locations of (1) the LIC binding region (2) the heavy chain binding region (3) the coiled-coil and (4) the spindly motif labelled. **C.** Schematics of the adaptors used in this study: BicD2 (blue), NINL (green), KASH5 (pink) and Hook3 (orange). The domains outlined in panel B are labeled on each schematic. Dashed lines indicate random coils in the protein. **D.** Single-molecule velocity (µm/sec) of activated dynein-dynactin-adaptor complexes: DDB (BicD2), DDN (NINL), DDK (KASH5) and DDH (Hook3). Error bars are median ± interquartile range. Statistical analysis was performed using a Kruskal-Wallis with Dunn’s multiple comparisons test. p-values and n can be found in Supplemental file 1. **E.** Single-molecule run lengths (µm) of DDB, DDN, DDK and DDH complexes. Error bars are median ± interquartile range. Statistical analysis was performed using a Kruskal-Wallis with Dunn’s multiple comparisons test. p-values and n can be found in Supplemental file 1. **F.** Landing rates of processive DDB, DDN, DDK and DDH complexes reported as events per micrometer of microtubule per nanomolar dynein per minute. Error bars are median ± interquartile range. Statistical analysis was performed using a Brown-Forsythe and Welch ANOVA with Dunnett’s T3 multiple comparisons test. p-values and n can be found in Supplemental file 1. **G.** Example kymographs of DDB, DDN, DDK and DDH. **H.** Schematic of the peroxisome relocalization assay: Pex3-GFP-FKBP and Halo-Adaptor-FRB dimerize upon addition of A/C Heterodimerizer, driving dynein-mediated clustering of peroxisomes in cells. **I.** Normalized peroxisome coverage at 0, 10 and 30 min in cells transfected with KASH5 or NINL. Error bars indicate mean ± SEM. Statistical analysis was performed using a Mann-Whitney test at each time point. n = 33-50 cells across three biological replicates. p-values can be found in Supplemental file 1.

In the absence of additional cofactors, dynein is in an autoinhibited conformation, called Phi, and cannot move processively on the microtubule (Zhang et al., 2017; Torisawa et al., 2014). To be active, dynein must bind to the multi-subunit protein complex dynactin and one of about 20 cargo adaptors (adaptors) (Reck-Peterson et al., 2018; Lee et al., 2020; Olenick and Holzbaur, 2019). Together, dynein, dynactin, and adaptors form the *active transport complex*, which converts dynein into an active conformation that is competent to move processively along the microtubule (Figure 1A) (McKenney et al., 2014; Schlager et al., 2014).

All known adaptors fall into one of four families, grouped according to the identity of a domain in their N-terminus that mediates binding to the light intermediate chain. These domains, which we will refer to as light intermediate chain binding (LIC-B) domains, include a coiled-coil (CC1-box family), an EF-Hand (EF-Hand family), a Hook domain (Hook family), or an RH domain (RH family) (Olenick and Holzbaur, 2019; Reck-Peterson et al., 2018; Singh et al., 2024). Despite having distinct structures, each type of LIC-B domain engages with the same short helix in the light intermediate chain, termed helix-1. In fact, X-ray crystal structures of helix-1 peptide bound to the LIC-B domain from CC1-box, EF-hand, and Hook type adaptors show that these LIC-B domains all form shallow hydrophobic pockets that associate with the same face of helix-1, burying two highly conserved phenylalanine residues (Lee et al., 2020, 2018).

In addition to a LIC-B domain, all adaptors also have a long, coiled-coil (CC) domain that is located on the C-terminal side of the LIC-B domain and interacts with dynein heavy chain and multiple dynactin subunits (Chaaban and Carter, 2022; Singh et al., 2024; Grotjahn et al., 2018; Urnavicius et al., 2015, 2018). Between adaptors, the CC domains show very little sequence conservation, except for two weakly conserved motifs that can be identified in many adaptors. The first motif is called the heavy chain binding site (HCBS) and mediates binding to heavy chain tails and a region of the intermediate chain (Chaaban and Carter, 2022). The second is called the spindly motif and mediates dynactin-specific interactions (Figure 1B) (d’Amico et al., 2022; Gama et al., 2017).

Adaptors are partially responsible for dynein’s ability to traffic cargo with specificity because different adaptors localize to different types of cargo (Reck-Peterson et al., 2018; Park et al., 2024; Olenick and Holzbaur, 2019). While all known adaptors have an LIC-B and a CC domain, adaptors are more divergent at their far C-terminal end and contain domains or sequences that facilitate cargo-specific interactions and localization (Park et al., 2024; Redwine et al., 2017). For example, four adaptors used in this study (Figure 1C) all link dynein to different cargos: BicD2 links dynein to Golgi, exocytic vesicles, and the nucleoporin RanBP2 (Gallisà-Suñé et al., 2023; McKenney et al., 2014; Hoogenraad et al., 2001; Matanis et al., 2002; Splinter et al., 2010); NINL may mediate Rab8-MICAL3 movement and link dynein to the centrosome (Redwine et al., 2017; Bachmann-Gagescu et al., 2015; Stevens et al., 2022); Hook3 links dynein to endosomes (Bielska et al., 2014; Schroeder and Vale, 2016; Bielska et al., 2014; Walenta et al., 2001); and KASH5 links dynein to meiotic chromosomes across the nuclear envelope (Agrawal et al., 2022; Horn et al., 2013; Garner et al., 2023).

Multiple high-resolution structures of dynein-dynactin-adaptor complexes have revealed key interfaces between all three components that are critical for stabilizing the final activated complex. For example, the adaptors’ CC domains bind dynein heavy chain tail and multiple subunits in dynactin, while the spindly motif specifically coordinates adaptor association with the pointed-end of dynactin (Chaaban and Carter, 2022; Singh et al., 2024; Urnavicius et al., 2018, 2015). However, existing cryo-EM structures don’t fully explain the importance of every interaction known to be essential for dynein activity. For example, despite being critical for dynein activity in cells and *in vitro*, the binding interface between dynein light intermediate chain helix-1 and the adaptors’ LIC-B is not well resolved in determined structures (Gama et al., 2017; Schroeder and Vale, 2016; Dwivedi et al., 2019; Chaaban and Carter, 2022; Singh et al., 2024; Urnavicius et al., 2018). This is likely partially due to the high flexibility of the light intermediate chain region that contains helix-1, but it is also possible that this interface is not maintained in the final complex. Indeed, though it is well established that disruption of the LIC-B – helix-1 interaction inhibits dynein activation, the molecular function of this interaction is unknown (Lee et al., 2020, 2018). For example, it is possible that light intermediate chain-adaptor binding is required to initiate complex formation but is not essential for persistent dynein movement after the complex is assembled.

Many groups have reconstituted active transport complexes with different adaptors and assessed the resultant motility. Generally, it seems that active transport complexes formed with different adaptors show differences in velocity, run length, or the ability to bind to the microtubule. (McKenney et al., 2014; Singh et al., 2024; Elshenawy et al., 2019; Htet et al., 2020). This is remarkable, as cryo-EM structures determined for active transport complexes formed with different adaptors don’t show large differences in dynein conformation (Singh et al., 2024; Chaaban and Carter, 2022; Grotjahn et al., 2018; Urnavicius et al., 2018). This suggests that any *bona fide* differences in dynein motility caused by adaptor identity is the result of either subtle structural changes in the final active structure or differences in the mechanism of how the active transport complex forms that would ultimately influence the amount of dynein that can achieve the active state.

The observation that distinct adaptors may form dynein-dynactin-adaptor complexes with divergent motility properties is interesting because, *in vivo* there are striking differences in the velocity, run length, or directional persistence of retrograde-directed cargo, even though these movements are all driven by dynein. For example, endosomes typically go on long, fast excursions (reaching velocities above 1 μm/sec), while meiotic chromosome movements driven by dynein are halting and slow (rarely reaching velocities above 100 nm/sec in mammals) (Flores-Rodriguez et al., 2011; Lee et al., 2015). One intriguing possibility is that different adaptors form active transport complexes with different motility behaviors that are tuned for the needs of the specific cargo that they interact with. If this is true, not only would it partially explain the diversity in retrograde trafficking behavior of different classes of cargo, but it would also begin to explain how the single dynein motor can be tuned to meet the unique trafficking requirements of divergent cargos.

To begin to explore if differences in the LIC-B and CC domain sequences of adaptors can alter dynein activity, we assessed the motility of active transport complexes formed with four canonical adaptors (BicD2, NINL, KASH5 and Hook3) using single-molecule TIRF microscopy. We found that different adaptors indeed modulate dynein’s motility properties such as velocity, landing rate and run length differently and that this directly leads to different cargo trafficking properties in cells. To understand the molecular features in adaptors that enable the observed differences, we assessed the activity of several truncated adaptor variants as well as a large library of chimeric adaptors that we generated by systematically swapping the LIC-B and CC domains of each canonical adaptor. We made several findings. First, we found that the CC domain is necessary and sufficient for dynein activation, though the LIC-B domain potentiates the dynein-activation ability of an adaptor. We also found that generally, the LIC-B and CC domains are modular and that any adaptor that has a LIC-B and CC domain can activate dynein motility above a certain threshold. Remarkably, we found that all canonical adaptors in this study could be engineered (via sequence alterations) to generate active transport complexes that show an increase in dynein’s velocity, run length, or landing rate. We speculate that this suggests that for most cargo, there is no selective pressure to generate adaptor sequences that can maximally activate dynein motility, and in fact, some adaptors sequences may have evolved to modestly activate dynein. Finally, we found that one of the engineered chimeric adaptors was hyperactive and could form active transport complexes with significantly higher fidelity than any canonical adaptor and supported faster cargo movement in cells. By characterizing this gain-of-function variant, we found that increased random coil at key positions in adaptor sequences facilitates robust and precise formation of the active transport complex. Our work supports a model where increased adaptor flexibility facilitates a kind of kinetic proofreading, where improperly assembled, inactive dynein-dynactin-adaptor complexes are specifically destabilized. Together, these results show that differences in adaptor domains can generate populations of transport complexes that display different motility. Further, this work suggests that specific features in adaptors could be harnessed to control or modulate cargo motility behavior in cells.

## Results

### Adaptors generate active transport complexes with distinct motility

The overarching motivation for this work was to identify the molecular features of adaptors that control their ability to activate dynein and explore if differences between adaptors have different effects on dynein motility. We selected four model adaptors from three canonical families: BicD2 (a member of the CC-box family), NINL (a member of the EF-hand family with two EF-hand pairs), KASH5 (a member of the EF-hand family with one EF-hand pair), and Hook3 (a member of the Hook family) (Figure 1C) (Reck-Peterson et al., 2018; Olenick and Holzbaur, 2019). In the absence of cargo, some adaptors assume an autoinhibited conformation where the C-terminal end folds back to interact with the N-terminal coiled-coil, thus blocking dynein and dynactin association (Splinter et al., 2010; Gibson et al., 2022). To avoid potential autoinhibition, we used well-characterized truncations of the adaptors that only contain the LIC-B and CC domains and have been previously shown to activate dynein *in vitro* (Htet et al., 2020; Agrawal et al., 2022).

To begin, we characterized the motility parameters (velocity, run length and landing rate) of active transport complexes formed with each adaptor. Though previous studies certainly support the hypothesis that many adaptors differ in their ability to activate dynein, we felt that it was important to establish the range of adaptor-dependent motility differences in side-by-side experiments performed with the same preparations of dynein and dynactin and in identical buffer conditions. To perform these experiments, purified dynein, dynactin, and adaptor were mixed at a 1:2:10 stoichiometric ratio and incubated for 20 minutes before dynein motility was assessed via TIRF microscopy in preassembled flow chambers containing Taxol-stabilized microtubules. In these experiments, we visualized dynein that was labeled with either JF646 or TMR linked via an N-terminal SNAP-tag. The 20-minute incubation is critical, as we previously observed that adaptors can form the active transport complex with different association kinetics (Gillies et al., 2025). As expected, adaptors formed complexes with different motility. While complexes formed with BicD2 and NINL were indistinguishable from each other with respect to all measured parameters, they were different from complexes formed with KASH5 and Hook3 (Figure 1D-G). KASH5 and Hook3 complexes varied from each other, as well. For example, BicD2, NINL, and Hook3 form complexes that move with similar velocities (∼600 nm/sec), while KASH5 complexes move with a much slower velocity (∼275 nm/sec) (Figure 1D). BicD2 and NINL complexes displayed the longest run lengths, KASH5 complexes had the shortest, while complexes formed with Hook3 had intermediate run lengths (Figure 1E). Finally, both KASH5 and Hook3 were generally less efficient at forming active transport complexes, displaying much fewer processive events (as determined by measuring the landing rate of only processive events) than BicD2 and NINL complexes (Figure 1F). These findings support the idea that active transport complexes formed with different adaptors fundamentally differ in their motility.

### Motility properties measured in vitro relate to cargo trafficking behavior in cells

Given that active transport complexes formed *in vitro* show striking differences in motility properties, we wondered if some of the observed differences in dynein-driven cargo movement in cells is caused by the differences in adaptor sequence. To test the possibility that the identity of the adaptor can influence the way dynein moves cellular cargos, we leveraged a well-established peroxisome relocalization assay (Figure 1H). Dynein-driven peroxisome trafficking, which is typically infrequent, can be induced by artificially recruiting adaptors to the organelle surface through the inducible FKBP/FRB heterodimerization system. This activates dynein on peroxisomes, leading to juxta-nuclear clustering. This approach allows us to directly compare the motility of the same cargo linked to dynein by different adaptors.

We set out to test if the model adaptors NINL and KASH5 differed in their ability to support dynein-driven peroxisome trafficking. We chose these two adaptors because they are both members of the EF-hand family, yet they differ dramatically in the motility properties that they impart to dynein *in vitro*. We assessed dynein-driven peroxisome trafficking by measuring relative peroxisome coverage (i.e. the total area of peroxisomes divided by the cell area) as a function of time after inducing adaptor recruitment to peroxisomes. Lower coverage values indicate more dynein-driven peroxisome clustering. Strikingly, we observed significant differences in the rate of dynein-driven peroxisome clustering between NINL and KASH5 (Figure 1I, S1A-B). At 10 minutes, peroxisomes in NINL-expressing cells were significantly closer to the nucleus than they were at the start of the experiment, and by 30 minutes nearly all peroxisome signal was confined to a bright punctum in the center of the cell (which is most likely the centrosome). In contrast, peroxisomes tethered to dynein by KASH5 clustered more slowly, and only displayed an increase in juxtanuclear localization 30 minutes after adaptor recruitment. These results show that cargo moved by dynein bound to NINL are trafficked more efficiently than those linked via KASH5. This finding is consistent with the *in vitro* observation that NINL generates active transport complexes that move faster, farther, and more often than those formed with KASH5. More generally, these results suggest that 1) *in vitro* motility experiments performed with different adaptors can inform predictions about cellular cargo movement; and 2) differences in adaptor identity likely contributes to the diversity in retrograde trafficking behavior observed for various cellular cargo.

### The adaptor coiled-coil is necessary and sufficient for promoting motility

Generally, adaptors can be segmented into two broad regions: the LIC-B and the CC domains (Figure 1B). We next asked if both domains were required for an adaptor to support the formation of an active transport complex. We chose NINL as the adaptor to modify for these experiments because it forms highly active dynein-dynactin-adaptor complexes and has fast dynein binding kinetics (Gillies et al., 2025). First, we generated two constructs: NINL^LIC-B^, which only contains the LIC-B domain, and NINL^CC^, which contains the rest of the truncated NINL sequence and should only bind to dynein heavy chain and dynactin (Figure 2A). We confirmed that NINL^CC^ did not have a cryptic helix-1 binding site using ColabFold (Figure S2A-B). Next, we asked if either construct could support dynein motility. We observed that, when mixed with dynein, dynactin, and Lis1 (a regulator that promotes dynein-dynactin-adaptor complex formation), NINL^CC^ could robustly activate dynein motility, albeit to a lesser extent than NINL (Figure 2B-C, S2C-D) (Htet et al., 2020; Elshenawy et al., 2020; Gutierrez et al., 2017; Markus et al., 2020). However, NINL^LIC-B^ did not activate motility on its own, nor did it increase the landing rate of complexes formed with NINL^CC^ when both constructs were included (Figure 2B-C, S2C-D). These data show that the CC domain is necessary and sufficient for dynein motility.

**Figure 2:**
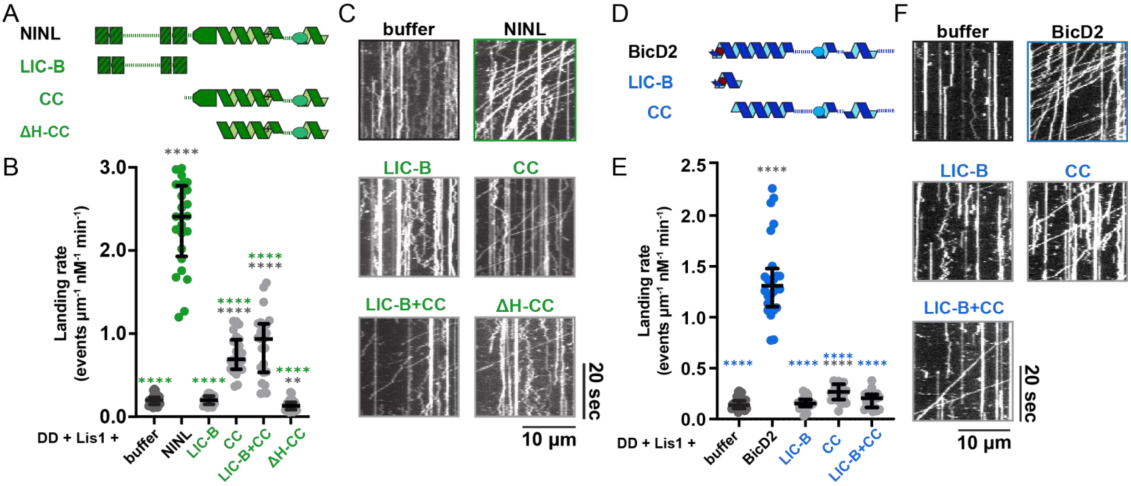
Adaptor coiled-coil is necessary and sufficient for promoting motility in the presence of Lis1. **A.** Schematic of the NINL adaptor and the different truncated constructs: NINL^LIC-B^, NINL^CC^, and NINL^ΔH-CC^. **B.** Landing rate of DD-Lis1 complexes incubated with either buffer (dark gray dots), NINL (green dots), NINL^LIC-B^, NINL^CC^, a combination of NINL^LIC-B^ and NINL^CC^, or NINL^ΔH-CC^, reported as processive events per micrometer of microtubule per nanomolar dynein per minute. Error bars are median ± interquartile range. Statistical analysis was performed using a Brown-Forsythe and Welch ANOVA with Dunnett’s T3 multiple comparisons test. Samples are compared to NINL (green stars) or buffer (dark gray stars). p-values and n can be found in Supplemental file 1. **C.** Representative kymographs of DD-Lis1 complexes incubated with either buffer, NINL, NINL^LIC-B^, NINL^CC^, a combination of NINL^LIC-B^ and NINL^CC^, or NINL^ΔH-CC^. **D.** Schematic of the BicD2 adaptor and the different truncated constructs: BicD2^LIC-B^ and BicD2^CC^. **E.** Landing rate of DD-Lis1 complexes incubated with either buffer (dark gray dots), BicD2 (blue dots), BicD2^LIC-B^, BicD2^CC^, or a combination of BicD2^LIC-B^ and BicD2^CC^ reported as processive events per micrometer of microtubule per nanomolar dynein per minute. Error bars are median ± interquartile range. Statistical analysis was performed using a Brown-Forsythe and Welch ANOVA with Dunnett’s T3 multiple comparisons test. Samples are compared to BicD2 (blue stars) or buffer (dark gray stars). p-values and n can be found in Supplemental file 1. **F.** Representative kymographs of DD-Lis1 complexes incubated with either buffer, BicD2, BicD2^LIC-B^, BicD2^CC^, or a combination of BicD2^LIC-B^ and BicD2^CC^.

ColabFold predictions of NINL’s structure show a small helical domain that caps the N-terminus of NINL’s CC (Figure S2A). To test if this domain is required for the activity of NINL^CC^, we generated a construct based on NINL^CC^ with this domain deleted and refer to this construct as NINL^ΔH-CC^ (Figure 2A). Unlike NINL^CC^, NINL^ΔH-CC^ did not activate dynein motility, which suggests that this domain either stabilizes NINL’s coiled-coil or participates in binding to dynein or dynactin (Figure 2B-C, S2C-D). Structural studies of dynein-dynactin-NINL complex will be required to resolve this question.

To test if the CC domains from other adaptors were also necessary and sufficient to facilitate dynein motility, we repeated the experiments outlined above with constructs generated from BicD2 (Figure 2D). As with NINL, we made constructs containing just the BicD2 LIC-B and CC domains and found that only the CC domain was needed to generate processive active transport complexes (Figure 2E-F, S2E-F). Collectively, these results suggest that the function of the LIC-B domain in all adaptors is to facilitate complex formation by binding to the light intermediate chain but is not required for sustained movement after the initial encounter complex forms.

### Chimeric adaptors reveal the breadth of plasticity in adaptor sequence

Our previous result illustrated that while the CC domain was required to form an active transport complex, the segment containing the LIC-B domain amplified the adaptors’ activation potential. To begin to dissect how each adaptor segment contributed to driving dynein activation, we generated “domain-swapped” chimeric adaptors, formed by linking the LIC-B domain of one adaptor to the CC domain of another. Using BicD2, NINL, KASH5, and Hook3 as “parents”, we made and purified each of the 12 possible swapped chimeras. Moving forward, we will refer to the domain-swapped chimeric adaptors with two capital letters, where the first letter indicates the identity of the parent adaptor from which the LIC-B domain originates, and the second letter indicates the identity of the parent from which the CC domain originates. For example, the chimera “BH” is generated by fusing the LIC-B region from BicD2 to the CC domain of Hook3, while “HB” is the chimera generated from the fusion of the LIC-B and CC domains of Hook3 and BicD2, respectively (Figure 3A).

**Figure 3:**
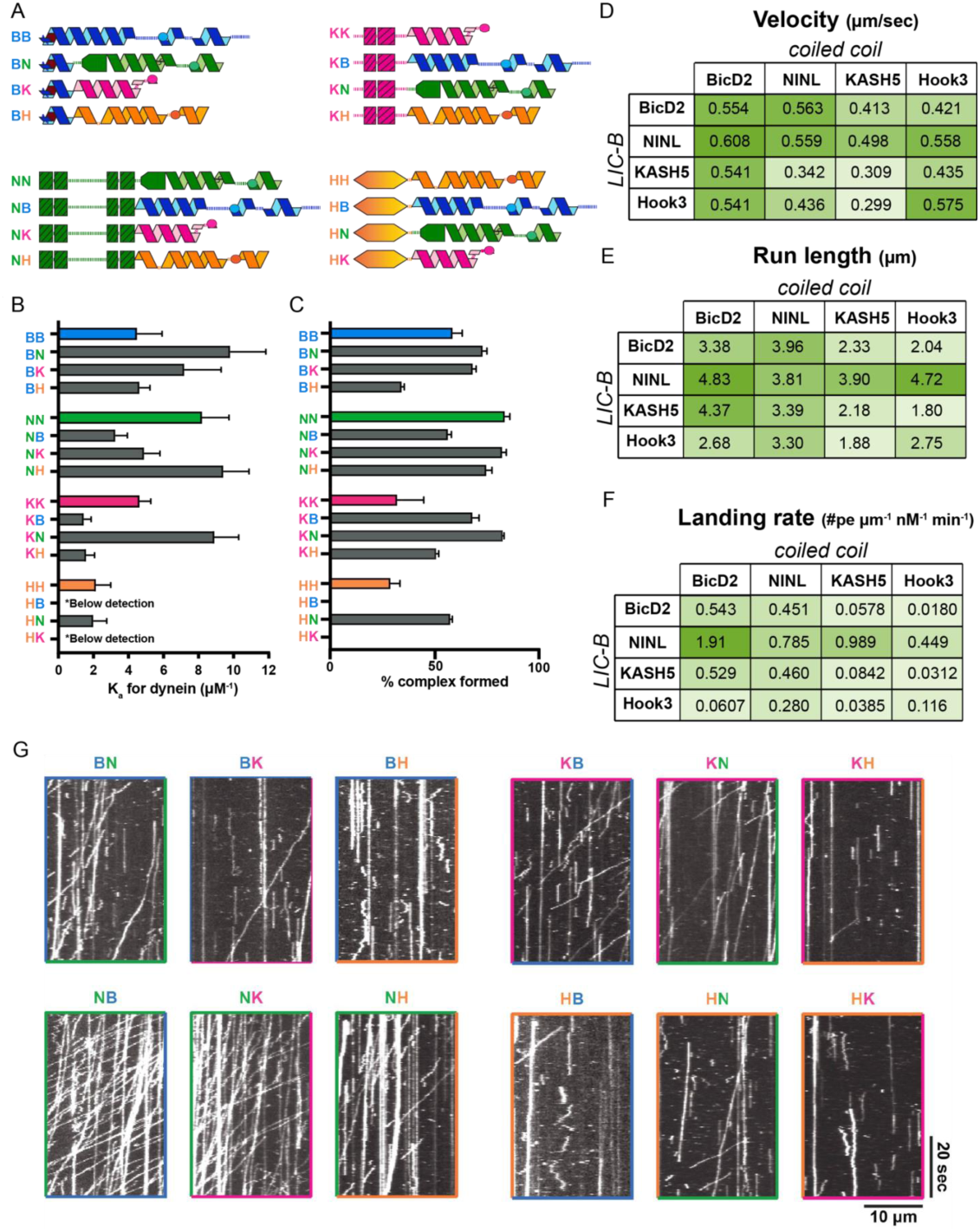
Chimeric adaptors activate dynein motility to varying degrees. **A.** Schematic of chimeric adaptors grouped by the identity of their LIC-B domains. The parent adaptors are shown at the top of each group. **B.** K_a_ (µM^-1^) of different adaptors (parent and chimeras) for dynein. Error bars indicate 95% confidence intervals. Raw data can be found in Figure S3.1. Data was collected from three independent experiments. **C.** Percentage of dynein-dynactin-adaptor complexes formed in the presence of different adaptors (parent and chimeras) as measured by dynein depletion in the presence of dynactin. Error bars indicate 95% confidence intervals. Data was collected from three independent experiments. **D-F.** Median single-molecule **(D)** velocity (µm/sec), **(E)** run length, and **(F)** landing rate (processive events/ µm nM min) of dynein complexes formed with different parent and chimeric adaptors are shown as tables, where each cell represents an adaptor. The identity of the LIC-B domain in an adaptor is indicated by the columns on the left, and the identity of the coiled-coil is indicated by the top rows. The range of values (velocity, run length or processive landing rate) observed with different adaptors is reflected by the color gradient in each table, where lighter green shades represent lower median values and darker green shades indicate higher median values. Raw data can be found in Figure S3.2. **G.** Example kymographs of dynein complexes formed with chimeric adaptors.

First, we simply asked if the chimeric adaptors were able to bind dynein using an *in vitro* quantitative pulldown assay. We observed that all but four (KB, HB, KH, and HK) of the chimeric adaptors could bind dynein with a K_a_ higher than 1.6 μM^-1^, which is the lower detection limit of the binding assay (Figure 3B, S3.1A-D). In fact, of the chimeras that bound robustly, none had K_a_ values for dynein that deviated more than 1.6-fold from one of the adaptor parents (Figure 3B). Interestingly, five chimeras (BN, BK, BH, KN, and NH) interacted with dynein with a higher affinity than both parents (Figure 3B, S3.1A-D).

Next, we asked if the chimeric adaptors deviated from their parents in their ability to form dynein-dynactin-adaptor complexes. To test this, we covalently linked each adaptor to a bead support and determined the percentage of dynein that is depleted by each adaptor in the presence of dynactin. Except for two adaptors (HB and HK), nearly all chimeras bound to dynein in the presence of dynactin as good or better than one or both parent adaptors (Figure 3C). We next asked if the ability to bind dynein in the absence of dynactin predicted the propensity to form dynein-dynactin-adaptor complexes by testing if the K_a_ values for dynein-adaptor binding (Figure 3B) correlated to the percent dynein bound in the presence of dynactin (Figure 3C) for each chimera.

Here (and in subsequent figures), data points from parent adaptors are solid-colored circles and chimeric adaptors are split color circles. The color on the left half of a chimera circle indicates the parent from with the LIC-B domain came and the color on the right indicates the origin of the CC domain. We observed a positive linear correlation between K_a_ and percent complex formed, which means that higher affinity dynein-adaptor binding is associated with increased ability to form dynein-dynactin-adaptor complexes (Figure S3.1E).

Next, we asked if the chimeric adaptors could activate dynein in the TIRF-based motility assay described above and measured the motility properties of active transport complexes made with each chimeric adaptor. Surprisingly, we found that every chimera could activate dynein motility, as measured by the ability to generate processive, motile events (Figure 3D-G, S3.2A-C). Only three of the chimeric adaptors (BH, KH and HK) showed landing rates below all the parent adaptors (Figure 3F-G). Additionally, all chimeras (including BH, KH, and HK) generated complexes that were sensitive to Lis1, showing increased velocity, landing rate, and run length in the presence of Lis1 (Figure S3.3A-G). These findings show that for an adaptor to simply activate dynein above a certain threshold, the identity of the LIC-B domain is not sensitive to the sequence of the CC, and vice versa.

### Motility parameters are partially correlated

In characterizing the activity of each parent and chimeric adaptor, we generated a large dataset of the velocity, run length and landing rate of dynein-dynactin-adaptor complexes formed with 16 different adaptors, both in the absence and presence of Lis1. In aggregate, these data offer a unique opportunity to understand how each motility parameter relates to each other and to learn more about the factors that control the observable behavior of a moving motor protein. We began by exploring the potential relationships between the velocity, run length, and landing rate parameters of moving dynein-dynactin-adaptor complexes assembled in the absence of Lis1 (original data in Figure 3D-G). To do this, we plotted the median values of each measured parameter versus every other and assessed linear and monotonic correlation (Figure 4A-C, S4A-C). We found that run length had a strong linear correlation with both velocity and landing rate, however velocity and landing rate did not correlate with each other (Figure 4A-C). These observations indicate that the observable run length of dynein-dynactin-adaptor complexes is likely controlled by the combined effect of the other two motility parameters. In other words, complexes that move fast are more likely to travel further on the microtubule and complexes that have a high affinity for the track (i.e. have a high landing rate) are less likely to fall off the microtubule. The same positive relationship between run-length and velocity has been observed for cargos bound to many active transport complexes (Soundararajan and Bullock, 2014).

**Figure 4:**
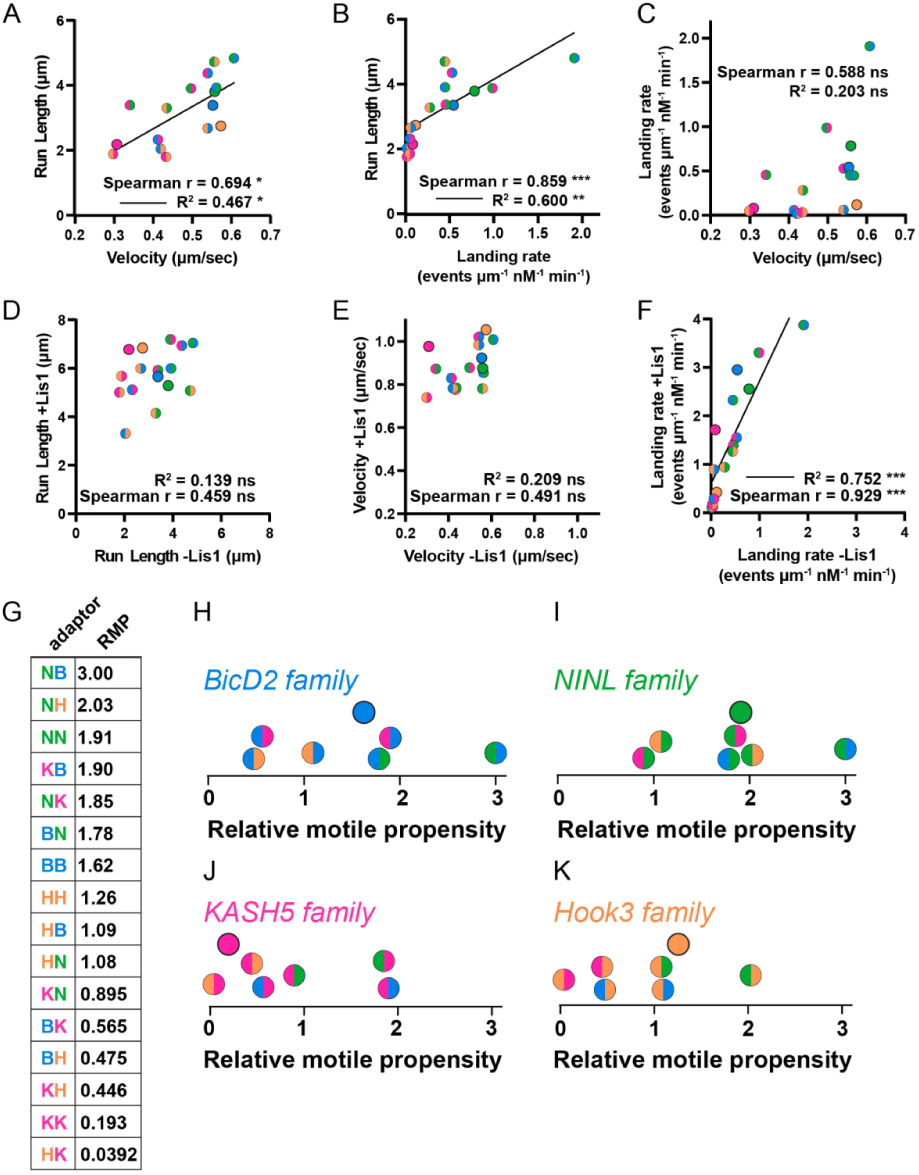
Motility parameters are partially correlated. **A-C.** Relation between **(A)** median run length (µm) and median velocity (µm/sec), **(B)** median run length (µm) and median landing rate (processive events/µm nM min), and **(C)** median landing rate (processive events/µm nM min) and median velocity (µm/sec) of active dynein complexes formed with different adaptors as measured in single-molecule motility assays. Each circle represents an adaptor, where the color in the first half of the circle represents the identity of the LIC-B domain and the color in the second half of the circle represents the identity of the coiled-coil domain. Spearman’s Correlation coefficient (r) and Pearson’s coefficient of determination (R^2^) are indicated. Solid line shows the linear fit to the data. p-values can be found in Supplemental file 1. **D-F.** Influence of Lis1 on different motility parameters. Relation between **(D)** median run length (µm), **(E)** median velocity (µm/sec), and **(F)** median landing rate (processive events/µm nM min) of active dynein complexes formed with different adaptors in the presence (y-axes) and absence (x-axes) of Lis1. Each circle represents an adaptor, where the color in the first half of the circle represents the identity of the LIC-B domain and the color in the second half of the circle represents the identity of the coiled-coil domain. Spearman’s Correlation coefficient (r) and Pearson’s coefficient of determination (R^2^) are indicated. Solid line shows a linear fit to the data. p-values can be found in Supplemental file 1. **G**. Table showing the relative motile propensity (RMP) of dynein complexes formed with different adaptors. **H-K.** RMP values of dynein complexes formed with different adaptors grouped by the four parent (**(H)** BicD2, **(I)** NINL, **(J)** KASH5 and **(K)** Hook3) families. Each chimeric adaptor, because it is a combination of two parent adaptors, is shown in both families.

At first glance, the fact that velocity and microtubule binding affinity are causally associated with run length could appear trivial or even tautological. However, the linked nature of these parameters is not necessarily observed for other motors. For example, mutational analysis performed with several kinesin-1 and -3 motors shows that the run length of kinesins does not strongly correlate with either velocity or landing rate (Figure S4D-G) (Soppina and Verhey, 2014a). The linked nature of run length with velocity and landing rate is also not borne out of intrinsic properties of the dynein motor. For example, yeast dynein is capable of processive motility in the absence of dynactin and an adaptor, which allows for the examination of dynein motor activity in the absence of dynactin and adaptor. In this minimal system, mutational analysis has shown that velocity and microtubule binding affinity (as estimated by unbinding rate) are, in fact, strongly anti-correlated (Cleary et al., 2014). In other words, in the absence of dynactin and adaptor, the increased ATP turnover that is required to drive faster velocity is associated with a reduced microtubule binding affinity and often results in reduced run length, presumably because the probability that both heads will release from the track is increased (Cleary et al., 2014; Cho et al., 2008). Collectively, these observations indicate that, in addition to stabilizing an active dynein conformation, dynactin and adaptors likely play a key role in increasing the coordination of motor domains in the active transport complex. This accounts for the unique ability of active transport complexes to be both fast and highly processive.

We anticipated that the propensity for an adaptor to assemble into an active transport complex (original data shown in Figure 3C) would predict certain motility parameters, since movement is contingent on complex formation. However, our analysis showed that percent complex formation did not significantly correlate with any individual motility parameter (Figure S4A-C). We were particularly surprised that percent complex did not correlate with processive landing rate given dynein’s inability to bind and move on the microtubule without interacting with dynactin and an adaptor (Figure S4B). This suggests that there are additional factors besides dynein-dynactin-adaptor binding affinity that dictate the observable motile behavior of active transport complexes.

Next, we asked if sensitivity to activation by Lis1 correlated to motility behavior in the absence of Lis1. Here, we compared the velocity, run length, and landing rate of active transport complexes formed in the absence and presence of Lis1 (original data in Figure 3D-F, S3.3A-C). The velocity and run length variables without and with Lis1 were not strongly correlated (Figure 4D-E). In other words, though all adaptors showed an increase in velocity and run length with Lis1, the velocity and run length of motile complexes in the absence of Lis1 was not predictive of the relative impact that inclusion of Lis1 would have on these parameters. Interestingly, landing rate measurements with and without Lis1 showed a robust linear correlation, which suggests that affinity for the microtubule track without Lis1 is a strong predictor of how Lis1 will affect this parameter (Figure 4F).

### Relative motile propensity of chimeric adaptors suggest that motility properties may be tuned for specific biological functions

Our next goal was to systematically examine how each chimera diverged from its parents to understand the relationship between adaptor sequence and resultant dynein motility. To simplify this analysis, we collapsed the velocity, run length, and landing rate variables into one aggregate variable that we termed *Relative Motile Propensity* (RMP). To derive RMP, we normalized all median values for each parameter such that the lowest measured value was zero and the highest measured value was one, then summed the parameter values for each adaptor. By this measure, motor complexes that move faster, further, and land on the microtubule with higher frequency will have a larger RMP (with a maximum possible value of 3), while complexes that move slower, shorter, and land less frequently will have a lower RMP (with a minimum possible value of 0) (Figure 4G).

Next, we visualized the RMP values of each adaptor family on the same plots (Figure 4H-K). A family is defined as a single parent adaptor and all chimeras (i.e. “children”) that contain either the LIC-B or the CC domain from that parent. All chimeras are present on two family plots because they are made from components of two parents. Visualizing the RMP data in this way revealed several interesting observations. First, all parent adaptors had at least one child chimera with a higher RMP value. In other words, all parent adaptors could be made to go faster, further, or land more frequently by modifying their sequence. The RMP for the BicD2 and NINL parents was near the median value for each respective family, which indicates that these adaptors are highly plastic and the RMP can be tuned both up and down by changing their domain compositions (Figure 4H, I).

In contrast, the KASH5 and Hook3 parents showed divergent relationships with their respective children. Within the KASH5 family, except for HK, all chimeras had higher RMP than the KASH5 parent (Figure 4J). This suggests that nearly every modification made to KASH5 will result in an adaptor variant that can generate active transport complexes that move faster, further, and/or more frequently. Within the Hook3 family, except for NH, each chimera had lower RMP than the Hook3 parent, which indicates that changes made to Hook3 nearly always result in variants that will make dynein move slower, shorter, and/or less often (Figure 4K). This suggests that the LIC-B and CC domains of Hook3 operate in a quasi-cooperative way while mediating the formation of dynein-dynactin complexes. In other words, for Hook3 functionality, the identity of the LIC-B domain and the sequence of the CC are partially linked.

### The chimera generated from NINL LIC-B and BicD2 CC generates highly motile active transport complexes in vitro and in cells

The family analysis also revealed that the NINL LIC-B and the BicD2 CC domains were the most likely to generate adaptors with elevated RMP, as the chimeras with the two highest RMP in each family had either the LIC-B from NINL or the CC from BicD2 (Figure 4H-K). In fact, NB, which is generated from the fusion of these two domains, had an RMP value of 3, which means that of all adaptors in this study, NB generated active transport complexes that moved the fastest, farthest, and landed on the track most often. Though significant, the increase in velocity and run length for NB compared to the other adaptors was relatively modest (Figure 3D, E). However, the increase in the landing rate of complexes formed with NB compared to other adaptors was striking (Figure 3F). The landing rate of NB complexes (1.91 μm^-1^nM^-1^min^-1^) was nearly 2-fold higher than the next best adaptor (NK, with a landing rate of 0.989 μm ^-1^nM^-1^min^-1^), 2.5-fold higher than NINL (at 0.785 μm ^-1^nM^-1^min^-1^), and 3.5-fold higher than BicD2 (at 0.543 μm ^-1^nM^-1^min^-1^) (Figure 3F). NB’s landing rate is even greater than the sum of the parental rates (i.e. 1.33 μm^-1^nM^-1^min^-1^), suggesting that the chimeric adaptor NB is truly displaying gain-of-function-type behavior with respect to the landing rate parameter.

We next set out to assess if the hyperactive motility observed for NB would also manifest in a more complex, cellular system. To do this, we turned again to artificial peroxisome trafficking assay and asked if peroxisomes linked to dynein by NB differed in their movement from those linked via NINL. Because we were interested in assessing NB’s dynamic gain-of-function, we made a few modifications to the assay. Rather than assessing peroxisome clustering in fixed cells, we imaged co-transfected cells live so that we could directly measure the motility properties of moving peroxisomes in cells expressing NINL or NB. For both adaptors, rapid, processive peroxisome movement could be seen ∼4 minutes after adaptors were recruited to the peroxisome surface. To determine if peroxisomes driven by dynein bound to NINL or NB displayed differences in motility, we assessed the number of motile peroxisomes per cell, as well as the velocity and observable run length of each moving peroxisome for both adaptors (Figure 5A-C, S5A-D). It is important to note that this cell imaging data was acquired on a single focal plane, and moving peroxisomes often disappeared because they moved vertically out of focus. This means that the run length parameter measured in this assay is the lower bound of possible run lengths and not an absolute measurement.

**Figure 5:**
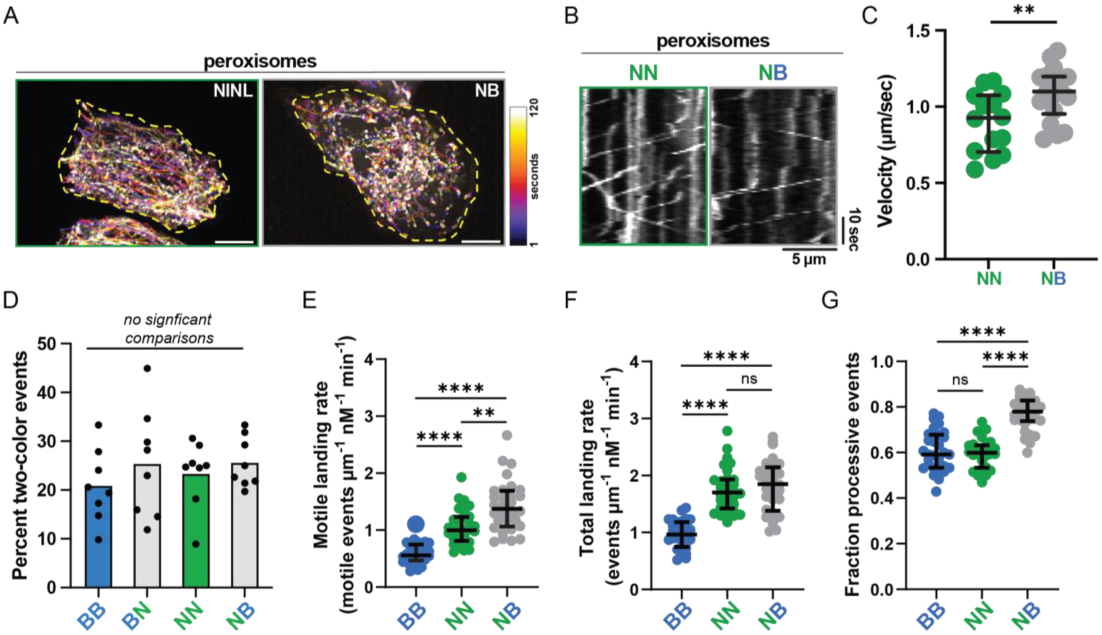
The NB super adaptor forms more motile-competent complexes *in vitro* and in cells. **A.** 2-minute temporal projection of fluorescently tagged peroxisomes moving in U2OS cells expressing NINL (left) or NB (right). Time scale indicates the start (1 sec - black) and end (120 sec - white) of the movie. Scale bar = 10µm. Yellow dashed line marks the cell boundary. **B.** Example kymographs showing the movement of peroxisomes in cells expressing NN (left) or NB (right). **C.** Velocity (µm/sec) of processively moving peroxisomes in U2OS cells expressing NN (green) or NB (gray). Error bars are median ± interquartile range. Statistical analysis was performed using a Mann-Whitney test. n= 14 cells for NINL and 18 cells for NB. p values can be found in Supplemental file 1. **D.** Percentage of two-color events in dynein-dynactin-adaptor complexes formed with BB, BN, NN and NB. p-values and n can be found in Supplemental file 1. **E.** Motile landing rate (processive events per micrometer of microtubule per nanomolar dynein per minute) for BB, NN and NB. Error bars indicate median ± interquartile range. Statistical analysis was performed using a Brown-Forsythe and Welch ANOVA with Dunnett’s T3 multiple comparisons test. p-values and n can be found in Supplemental file 1. **F.** Total landing rate (total events per micrometer of microtubule per nanomolar dynein per minute) for BB, NN and NB. Error bars indicate median ± interquartile range. Statistical analysis was performed using a Brown-Forsythe and Welch ANOVA with Dunnett’s T3 multiple comparisons test. p-values and n can be found in Supplemental file 1. **G.** Fraction of processive events (obtained as the ratio of processive to total events) for BB, NN and NB. Error bars indicate median ± interquartile range. Statistical analysis was performed using a Brown-Forsythe and Welch ANOVA with Dunnett’s T3 multiple comparisons test. p-values and n can be found in Supplemental file 1.

We anticipated that peroxisomes in cells expressing NB would move faster, longer, and more often than in cells expressing NINL since NB generated active complexes with a higher velocity, run length, and landing rate *in vitro*. Contrary to our expectations, there was no difference in the number of moving peroxisomes per unit cell area or the observable run length between each condition (Figure S5A-B). However, the prediction we made for velocity did bear out and we found that peroxisomes moved significantly faster in the NB condition, with median velocities ∼19% faster for cells expressing NB versus NINL (Figure 5C). Potential differences in expression level between the two adaptors does not explain this effect, as we observe no correlation between adaptor expression level and velocity (Figure S5C-D). Interestingly, the velocity increase observed in the peroxisome relocalization assay is larger than what we observed for NB in the fully reconstituted motility assay, where NB yields dynein that moves ∼9% faster than NINL. Together, these results confirms that the motility differences imparted to dynein by different adaptors *in vitro* translates to significant differences in cellular cargo trafficking and show that it is possible to engineer an adaptor to change cargo trafficking properties. However, this result also shows that the relationship between dynein motility and resultant cargo movement is complex. For example, an increased propensity for an adaptor to form more processive complexes *in vitro* does not simply translate into more cargo movement overall but instead likely supports faster cargo trafficking. More work will be required to establish all the factors that ultimately control the observable behavior of dynein-driven cargo movement and to establish how the behaviors observed *in vitro* relate to cargo movement in cells.

### Adaptors with increased flexibility are more likely to form motile active transport complexes

Why does NB generate more active dynein-dynactin-adaptor complexes? The most parsimonious explanation would be if NB has an elevated affinity for dynein or an increased propensity to bind dynein-dynactin. However, this does not seem to be the case. For example, we have already determined that NB’s affinity for dynein and its propensity to form dynein-dynactin-adaptor complexes in solution is comparable to BicD2 and lower than both NINL and its sibling BN (Figure 3B-C).

Previous studies have shown that adaptors often form transport complexes that contain two dynein dimers and that these complexes move faster than those formed with a single dynein dimer (Urnavicius et al., 2018). We wondered if the increased propensity for NB to form more motile complexes could be because it enabled the formation of more two dynein dimer complexes. To assess this, we repeated the motility experiments performed with NINL, BicD2, NB, and BN but included two different batches of dynein that were labeled with different fluorophores and determined the percentage of events that showed two-color dynein colocalization (as these represent unambiguous complexes with two dynein dimers). We observed no difference in the fraction of two-color dynein events for any adaptor in this family, suggesting that NB’s remarkable ability to activate dynein does not stem from its ability to control the stoichiometry of dynein motors in the transport complex (Figure 5D).

In every dynein motility experiment, a fraction of the dynein bound to the microtubule appear immotile. Dynactin and adaptor are nearly always colocalized with these “dead” events (Figure S5E), which shows that these events represent dynein-dynactin-adaptor complexes that are not capable of processive motility (Redwine et al., 2017). We wondered if NB could be displaying elevated activity because it was more efficient than other adaptors at forming *active* dynein-dynactin-adaptor complexes. In other words, we speculated that NB may form less inactive dynein-dynactin-adaptor complexes, which would not necessarily be reflected in bulk measurements of dynein-dynactin-adaptor affinity (as measured in Figure 3C). Because we previously only measured the landing rate of processive events, the data we have presented thus far did not allow us to assess the percent of dynein-dynactin-adaptor complexes that were active. To perform this measurement, we repeated the motility experiments as described previously but in addition to assessing landing rate of processive events, we also measured total events (including dead and diffusive) and calculated the percent of total events that were processive. In this series of experiments, we conjugated a fluorophore to each adaptor and restricted our analysis to events that were positive for adaptor signal. This ensured that measured immotile events were not simply free dynein (i.e. dynein not in the active transport complex). We performed these experiments with BicD2, NINL, and NB (Figure 5E-G, S5F).

First, we compared the parent adaptors BicD2 and NINL to each other. NINL showed a greater number of total events and more processive events compared to BicD2 (Figure 5E-F). However, the fraction of processive events (calculated by dividing number of processive events by total events) was the same for both parents (Figure 5G). This means that while NINL may form more complexes overall than BicD2, these complexes are not likely to be more active than those formed with BicD2. In contrast, we found that the total number of dynein-dynactin-NB events was not significantly different from NINL, but that the number of motile events was higher (Figure 5E-F). This means that NB has a significantly higher fractional processivity than either parent (Figure 5G). This result indicates that dynein-dynactin-adaptor complexes formed with NB are more likely to be motile and processive and shows that NB doesn’t simply form more complexes overall. In other words, NB forms active transport complexes with higher fidelity than either of its parents.

Our findings thus far show that NB can generate higher percentage of active transport complexes that land and move on the microtubule. We next asked what molecular feature in NB facilitated this ability. To begin, we compared the ColabFold structural predictions of BicD2, NINL, and their children chimeras BN and NB (Figure 6A, S6A-D). The most striking difference we observed was that NB’s sequence contains the highest percentage of random coil compared to other members of the family because it is generated by the fusion of the most flexible domains from both parents: NINL’s LIC-B contains two EF-hand pairs separated by ∼120 amino acids of random coil and the BicD2 CC domain contains ∼65 amino acids of random coil immediately before the spindly motif.

**Figure 6:**
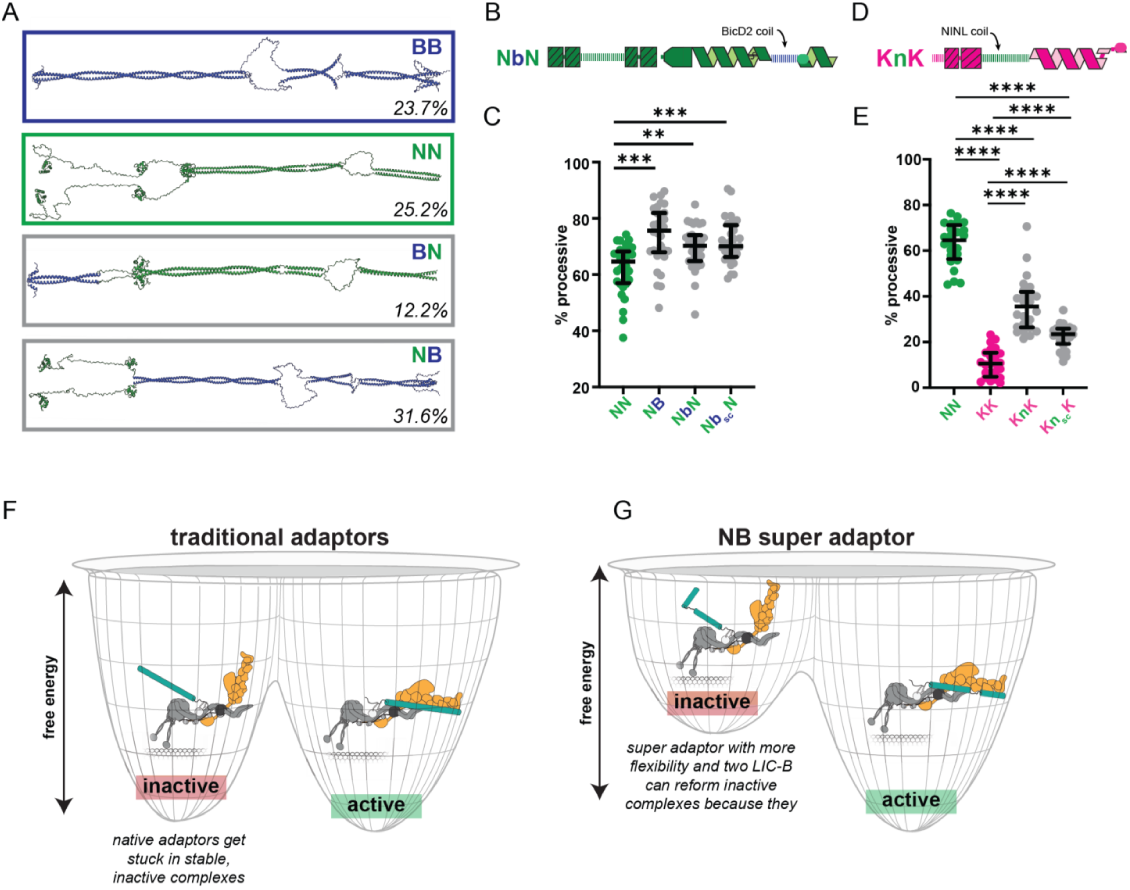
Random coils in the adaptor can improve dynein motility. **A.** ColabFold predictions of the NB family of adaptors- BicD2, NINL, BN and NB. The color of the domains indicates which parent they originate from (BicD2- blue; NINL- green). Flexible regions (with low reliability) in these structures were artificially ‘linearized’ using *Coot* for better visualization. The percentage of random coil in the protein is indicated in the bottom right for each adaptor. The PAE plots for these predicted structures can be found in Figure S6 A-D. **B-C**. **(B)** Schematic of the NbN construct and **(C)** the percent processivity (obtained as the percentage of processive to total events) for NN (green), NB, NbN and Nb_sc_N. Error bars indicate median ± interquartile range. Statistical analysis was performed using a Brown-Forsythe and Welch ANOVA with Dunnett’s T3 multiple comparisons test. p-values and n can be found in Supplemental file 1. **D-E**. **(D)** Schematic of the KnK construct and **(E)** the percent processivity (obtained as the percentage of processive to total events) for NN (green), KK (pink), KnK and KN_sc_K. Error bars indicate median ± interquartile range. Statistical analysis was performed using a Brown-Forsythe and Welch ANOVA with Dunnett’s T3 multiple comparisons test. p-values and n can be found in Supplemental file 1. **F-G.** Cartoon illustrating our proposed model for how dynein complexes assemble in the presence of either **(F)** traditional adaptors or **(G)** the NB super adaptor.

We hypothesized that increasing the random coil content may generate adaptors that form active transport complexes with increased motility. To test this hypothesis, we replaced the short stretch of random coil preceding the spindly motif in NINL (amino acids 579-614), with the corresponding residues from BicD2 (amino acids 269-334). We refer to this variant as NbN, where the lowercase “b” is designating random coil from BicD2 (Figure 6B, S6E). We next asked if NbN performed like NB in *in vitro* motility assays, and assessed the velocity, run length, processive landing rate, and percent processivity of dynein-dynactin-NbN complexes (Figure 6C, S6F-H). Interestingly, we found that NbN did perform better in both the run length and velocity parameters than the NINL parent (Figure S6F-G). NbN outperformed NINL in velocity, landing rate, and percent processivity, though it did not reach the same elevated level of N (Figure 6C, S6F-H). This result shows that the random coil from BicD2’s CC domain alone can confer increased activity to adaptors. We next wondered if the specific amino acid sequence or the length of the BicD2 CC random coil resulted in the increased functionality. To differentiate these possibilities, we generated another NINL-based variant that contained a stretch of random coil sequence generated by randomly reordering the amino acids that make up BicD2’s original CC random coil (Nb_sc_N, where b_sc_ indicates a scrambled sequence of the amino acids that make up the random coil in BicD2’s CC). Structural predictions confirmed that the scrambled sequence did not have a high propensity to assume unexpected secondary structure. We found that Nb_sc_N performed very similarly to NbN (Figure 6C, S6F-H), suggesting that the length of the coiled-coil segment, rather than the ordered sequence, is responsible for the increased activity.

We next asked if random coil in the LIC-B domain also contributed to the increased activity seen with NB. Rather than delete random coil from NB and potentially observe a decrease in activity, which would be challenging to interpret, we opted to introduce random coil into an adaptor with a lower RMP to test if we could increase its ability to activate dynein. To accomplish this, we replaced amino acids 121-154 in KASH5 with amino acids 81-196 from NINL. This construct, KnK contains KASH5’s EF-hands linked to KASH5 CC-domain with the random coil found in NINL’s LIC-B (designated by the lower case “n”) (Figure 6D, S6I-L). We also generated this same construct except with the NINL random coil sequence scrambled (Kn_sc_K) to differentiate the effect of amino acid sequence from coil length, as described above. Interestingly, KnK and Kn_sc_K both formed complexes that moved with an elevated velocity, run length, landing rate and percent processivity as compared to KASH5 (Figure 6E, S6M-O). These data show that increasing the flexibility of the linkage between the LIC-B and CC-domains by adding random coil sequence also increases an adaptors’ ability to form active transport complexes that move faster, more processively, and land more often on the microtubule track.

## Discussion

We set out to assess the possibility that different adaptors could give rise to active transport complexes with distinct motility and to understand if specific molecular features in adaptors controlled how the adaptors activate dynein. We found that between four canonical adaptors, BicD2, NINL, KASH5, and Hook3, there were large and striking differences in the observed motility as measured by assessing velocity, run length, and landing rate of moving complexes formed with each adaptor, highlighting the tremendous functional plasticity inherent in the dynein motor. We also found that the differences in motility measured *in vitro* predicted the efficiency with which different adaptors support cargo trafficking in cells. These finding suggests that different adaptors generate dynein-dynactin-adaptor complexes that move differently and that adaptor identity could underlie some of the variety of retrograde cargo trafficking behaviors observed in cellular settings.

By generating constructs that separated the LIC-B from the CC domain for two adaptors, we also found that the CC-domain is necessary and sufficient for dynein activation, but that a contiguous LIC-B domain functions to strongly potentiate the ability of the CC domain to activate dynein motility. This finding prompted us to ask if the LIC-B and CC domains within adaptors functioned in an independent or modular way, or if instead, specific LIC-B domains could only function with specific CC-domains to drive dynein activation. We generated a library of 12 synthetic adaptors, made by systematically swapping the LIC-B and CC domains between the four canonical model adaptors. To simply activate dynein, we found that nearly any LIC-B could operate with any CC domain, which shows that there is no fixed requirement for a given LIC-B to be linked to a specific CC domain.

In characterizing each chimera, we generated large datasets that gave us an opportunity to ask to if the motility parameters of velocity, run length, and landing rate were related to each other. In other words, we were able to ask what (if any) of the motility parameters control the others. We found that run length correlated strongly with velocity and landing rate, but that no other parameter or measurement correlated. This finding suggests that run length is controlled by velocity and microtubule binding affinity and, unsurprisingly, long, dynein driven excursions will result from fast moving complexes that have a high microtubule binding affinity. This trait is not shared with kinesins or with dynein from yeast that is active without dynactin and adaptor (Cleary et al., 2014; Soppina and Verhey, 2014a). Additionally, the causal relationship between velocity and microtubule binding affinity with run length does not emerge when multiple dyneins are linked together via artificial cargo in the absence of dynactin and adaptors (Driller-Colangelo et al., 2016) Together, these results indicates that dynactin and adaptors, in addition to overcoming dynein autoinhibition, function to increase motor-to-motor coordination within the active transport complex to ensure that all microtubule binding domains rarely enter the low affinity state at the same time.

While all chimeras could activate dynein, they did show differences in their activity. We assessed each chimera’s Relative Motile Propensity (RMP), which is an aggregate variable with contributions from velocity, run length, and landing rate. Chimeras across and within families showed broad RMP values, with some variants showing higher RMP values than the parent chimeras from which they were formed, while others had lower RMP. Interestingly nearly all modifications made to KASH5 generated a chimera variant that had an elevated RMP. In cells, KASH5 mediates the trafficking of chromosomes during prophase 1 of meiosis (Horn et al., 2013). The observed dynein-driven movement of these chromosomes is slow compared to other dynein cargos (with speeds of ∼50 - 100 nm/sec) and saltatory (Lee et al., 2015). It is likely that excessive or rapid dynein motility during meiosis could have adverse reactions on chromosome integrity or subsequent pairing and that the KASH5 adaptor sequence has specifically evolved to generate active dynein that moves with a low RMP. Future studies will be devoted to testing this hypothesis. Future studies will be devoted to testing this hypothesis. In contrast we found that some adaptors, like Hook3, are intolerant of alterations and nearly every chimera generated with Hook3 parts had lower RMP values than Hook3. These findings suggest that the identity of the Hook3 LIC-B domain may optimally function with the Hook3 CC domain and that Hook3, specifically, is less modular than the other adaptors.

Remarkably, one of the chimeric adaptors, NB, was a super adaptor and generated active transport complexes that moved faster, farther, and more often than every other adaptor tested. NB also supported faster peroxisome trafficking in cells than its NINL parent. We found that the increase in activity was because NB forms active dynein-dynactin-adaptor complexes with higher fidelity than other adaptors. In other words, NB doesn’t bind dynein and dynactin with a higher affinity than other adaptors, but dynein-dynactin-adaptor complexes formed with NB are more likely to be active than those assembled with other adaptors. This result shows that bulk affinity measurements of a given dynein-dynactin-adaptor complex can not necessarily predict the fidelity with which an adaptor will activate dynein. NB had the highest percentage of random coil of any adaptor in the library, and the structure-function studies we conducted confirmed that increasing the amount of random coil in an adaptor increases the ability for any adaptor to generate active dynein complexes with higher RMP values. This finding suggests that increased adaptor flexibility is associated with more motile dynein-dynactin-adaptor complexes.

Why is NB more likely to form active dynein-dynactin-adaptor complexes than other adaptors? How does increased random coil in the adaptor sequence increase the fidelity with which the adaptor can activate dynein? We have previously determined that complexes of dynein-dynactin-adaptor, once formed, are incredibly stable and rarely dissociate spontaneously (Gillies et al., 2025). While many of these complexes place dynein in an active conformation, our data shows that many dynein-dynactin-adaptor complexes, though long-lived, are not motile. For example, only about 60% of the total dynein-dynactin-adaptor complexes formed with NINL or BicD2 are capable of processive movement (Figure 5G). Inactive dynein-dynactin-adaptor species represent an off-pathway product that is improperly assembled such that dynein is not in an active conformation. Because both motile and immotile events are stable and long-lived (Figure S5G), we speculate that the free energy associated with inactive and active complexes formed with traditional adaptors is comparable. This means that a proportion of the dynein gets trapped in an inactive state (Figure 6F). For NB, we posit that the free energy associated with off-pathway, improperly assembled dynein-dynactin-NB complexes is less, which results in fewer dynein molecules trapped in the inactive conformation (Figure 6G). We reason that increased random coil in the adaptor sequence allows more conformational flexibility that functions to prevent NB from being trapped in improperly assembled complexes (Figure 6G). The model we postulate for the increased activity of NB is a type of kinetic proofreading, whereby improperly assembled complexes formed with NB dissociate rapidly, and get another chance to assemble correctly. Our previous work (Gillies et al., 2025) has shown that dynein-dynactin do not stably associate in the absence of an adaptor, which explains how simply increasing the ability for an adaptor to dissociate by increasing its flexibility could prevent dynein from being trapped in a local energy well associated with improper complex structure. Future studies will be focused on testing this model by monitoring the kinetics of NB binding to dynein and dynactin and by assessing the stability of motile and immotile complexes formed with NB and other adaptors.

## Contributions

M.E.D., J.P.G., and A.S. conceived of and planned the project. J.P.G., A.S., S.R.G., R.E.J., R.M., R.S.C., and C.C., generated reagents, collected data, and/or analyzed data for motility experiments. A.S., A.D., D.G., and J.L.Z. collected data, analyzed data, and/or generated analysis code for U2OS-based experiments. M.E.D. wrote the manuscript with input from all authors.

## Acknowledgements

The authors thank all members of the DeSantis, Cianfrocco, Ohi, Verhey, and Sept labs for helpful discussions. The authors also thank Drs. Richard Baker, John Salogiannis, and Bret Redwine for insightful feedback. This work was supported by NIH-R35GM146739, NSF-2142670, and NIH-R01HD108809 (to M.E.D), NIH-T32 GM145304 (to J.L.Z.), and NIH-T32GM132046 (to A.D).

## Methods

### Cloning and plasmid construction

The wild-type Lis1 construct (pFastBac backbone, for insect cell expression), along with the constitutively active adaptor constructs Hook3(1-552), NINL(1-702), and BicD2(1-598) in the pET28a backbone for bacterial expression, were a kind gift from the Reck-Peterson lab (UCSD). The stable HEK293 cell line expressing p62-Halo, used for dynactin purification, was also obtained from the Reck-Peterson lab. The dynein construct for insect cell expression (Addgene plasmid #111903) was kindly provided by the Carter lab (MRC). Constitutively active KASH5(1-400) was PCR-amplified using Q5 High-Fidelity 2X Master Mix (M0492L, NEB) and integrated into the pET28a backbone by isothermal assembly (Gibson Assembly Master Mix, E2611S, NEB), following the protocol described in (Gibson et al., 2009).

Chimeric adaptor constructs used in this study were generated by systematically exchanging the LIC-B and CC-domains from different adaptors. Domain boundaries were defined as follows: NINL (LIC-B: 1-275; CC: 276-702), BicD2 (LIC-B:1-98; CC: 58-598), KASH5 (LIC-B: 1-154; CC: 155-400), Hook3 (LIC-B: 1-160; CC: 161-552), where the numbers represent amino acid positions. Domain fragments were PCR-amplified using Q5 High-Fidelity 2X Master Mix (M0492L, NEB) and assembled into the pET28a backbone by isothermal assembly (Gibson Assembly Master Mix, E2611S, NEB), following the protocol described in (Gibson et al., 2009).

Truncated constructs of NINL (EF: 1-275; CC: 276-702; ΔH CC: 374-702) and BicD2 (CC1: 1-98; CC: 58-598) were generated by inverse PCR (Q5 High-Fidelity 2X Master Mix, M0492L, NEB) on the respective NINL and BicD2 parent adaptor plasmids, with primers flanking the region to be removed. The PCR product was then circularized using KLD Enzyme mix (M0554S, NEB) according to manufacturer’s instructions.

The random coil regions in NBscN (BicD2: 269-334) and KNscK (NINL: 81-196) were scrambled randomly and generated as DNA oligos (IDT) before being assembled with the respective NINL or KASH5 fragments.

All constructs used in this study were transformed into chemically competent DH5α cells by heat shock at 42 °C for 40 sec, followed by recovery in S.O.C media (Thermo Fisher Scientific) in a 37 °C shaker for 1 h before plating them on LB agar plates containing the appropriate antibiotic. The plates were incubated overnight at 37 °C and colonies were screened for the presence of the gene of interest by colony PCR. Positive colonies from the screen were cultured in LB media to isolate the plasmid, and the sequence was confirmed by sanger sequencing. A list of all constructs used in this study can be found in Supplemental File 1.

### Protein expression and purification

#### Dynein and Lis1 expression

Sf9 cells cultured in Sf-900 serum- free medium (Gibco) were used to express dynein and Lis1 as described (Garrott et al., 2023; Agrawal et al., 2022). Briefly, genes encoding human dynein carried in the pACEBac1 plasmid or wildtype Lis1 carried in pFastBac plasmid were transformed into competent DH10EmBacY cells *via* 42 °C heat shock for 15 s followed by a 6 h recovery in S.O.*C. media* (Thermo Fisher Scientific) at 37 °C with shaking at 220 rpm. Cells were then plated on LB-agar containing tetracycline (10 μg/ml), gentamicin (7 μg/ml), kanamycin (50 μg/ml), IPTG (40 μg/ml) and BluoGal (100 μg/ml). Blue/white screening was performed 48 to 72h after growth to identify colonies containing the plasmid of interest. For dynein, white colonies were selected and tested for the presence of all dynein genes by PCR. Selected colonies were then grown in LB media with tetracycline (10 μg/ml), gentamicin (7 μg/ml), kanamycin (50 μg/ml) in a 37 °C shaker at 220 rpm. After 14 to 16 h of incubation, bacmid DNA was isolated from cultures using isopropanol extraction. About 1 × 10^6^ Sf9 cells, cultured in six-well dishes, were subsequently transfected with 1 to 2 μg of the purified bacmid using FuGene HD transfection reagent (Promega), according to manufacturer’s instructions. Transfected Sf9 cells were grown in a humid incubator for 3 days at 27 °C without agitation. 1 ml of viral supernatant (designated V0) was then collected from the transfected wells by centrifuging at 1000*g*, for 5 min at 4 °C. The V0 stock was then used to infect a 50 ml culture of Sf9 cells at a density of 1 × 10^6^ cells/ml. The infected cells were incubated for 3 days in a 27 °C shaker at 105 rpm, after which the supernatant (designated V1) was harvested by centrifugation. For protein expression, 4 ml of the V1 stock was used to infect a 400 ml culture of Sf9 cells at a density of 1 × 10^6^ cells/ml. The culture was incubated again for 3 days in the 27 °C shaker at 105 rpm. Finally, the Sf9 cells were pelleted by centrifugation at 3500 rpm, washed with 10 ml of cold PBS, and flash-frozen in liquid nitrogen for storage at −80 °C until subsequently used for protein purification.

#### Dynein purification

Purification of human dynein was performed as described (Zang et al., 2025; Garrott et al., 2023). All purification steps were carried out at 4 °C unless indicated otherwise. Dynein pellets were thawed on ice and resuspended in 40 ml of dynein-lysis buffer per pellet (50 mM HEPES (pH 7.4), 100 mM sodium chloride, 1 mM DTT, 0.1 mM Mg-ATP, 10% (v/v) glycerol) supplemented with either 0.5 mM Pefabloc (Millipore-Sigma) and 1x cOmplete EDTA-free protease inhibitor cocktail (Roche) or 0.2mM AEBSF (Gold Bio), 81uM Bestatin (VWR), 7uM E-64 (Gold Bio), 21uM Leupeptin (Gold Bio), 7uM Pepstatin-A (VWR), 1mM Benzamidine (Millipore-Sigma), and 1mM PMSF (Millipore-Sigma). Cell lysis was achieved by using a Dounce homogenizer and the lysate was centrifuged (183,960*g*, 88 min, 4 °C) in a Type 70Ti rotor (Beckman) to obtain a clarified supernatant. 2 ml of IgG Sepharose 6 Fast Flow beads (Cytiva) were equilibrated with dynein-lysis buffer and then incubated with the clarified supernatant for 4 h with rotation to facilitate bead binding. A gravity column was then used to collect the beads, followed by washing with 200 ml of dynein-lysis buffer and 300 ml of tobacco etch virus (TEV) buffer (50 mM Tris–HCl (pH 8.0), 250 mM potassium acetate, 2 mM magnesium acetate, 1 mM EGTA, 1 mM DTT, 0.1 mM Mg-ATP, 10% (v/v) glycerol). For fluorescent labeling of the SNAP tag on dynein, dynein-bound beads were mixed with 5 μM SNAP-Cell-TMR-Star (NEB) or SNAP-JF646 (Lavis lab, Janelia), or SNAP-Surface Alexa 488 (NEB) for 10 min at room temperature. Beads were washed again with 300 ml of TEV-buffer, resuspended in 15 ml TEV buffer (supplemented with 0.5 mM Pefabloc and about 0.2 mg/ml TEV protease) and incubated overnight with gentle rotation. This mixture was passed through a gravity column to separate the cleaved proteins (in the flow through) from the beads. The flow through was then concentrated to 500 μl with a 100K MWCO concentrator (EMD Millipore), before injecting onto a TSKgel G4000SWXL size-exclusion column (Tosoh Bioscience) equilibrated with GF150 buffer (25 mM HEPES (pH 7.4), 150 mM KCl, 1 mM MgCl_2_, 1 mM DTT) supplemented with 0.1 mM Mg-ATP. The column was run at a flow rate of 0.75 ml/min and peak fractions were collected. These fractions were then buffer exchanged into GF150 buffer supplemented with 0.1 mg/ml ATP and 10% glycerol, concentrated to 0.1 to 0.5 mg/ml, flash frozen in small aliquots, and stored at −80 °C.

#### Lis1 purification

Lis1 was purified from Sf9 cells similarly, except the Lis1-lysis buffer (30 mM HEPES pH 7.4, 50 mM potassium acetate, 2 mM magnesium acetate, 1 mM EGTA, 300mM potassium chloride, 1 mM DTT, 0.5 mM Pefabloc and 10% (v/v) glycerol, supplemented with cOmplete EDTA-free protease inhibitor cocktail tablets (Roche)) was utilized in place of the dynein-lysis buffer. After lysis, centrifugation and bead binding, the mixture was passed through a gravity column and washed with 20 ml of Lis1-lysis buffer and 200 ml of Lis1-TEV buffer (10 mM Tris-HCl (pH 8.0), 2 mM magnesium acetate, 150 mM potassium acetate, 1 mM EGTA, 1 mM DTT, 10% (v/v) glycerol), supplemented with 100 mM potassium acetate and 0.5 mM Pefabloc, and finally with 100 ml of Lis1 - TEV buffer before incubating with 0.2 mg/ml of TEV protease overnight. The mixture was passed through a gravity column to collect the cleaved proteins, which were then concentrated to 500 μl with a 30K MWCO concentrator (EMD Millipore). The protein was injected onto a Superose 6 Increase 10/300 GL column (Cytiva) at 0.75 ml/min with GF150 buffer, supplemented with 10% (v/v) glycerol as the mobile phase. Peak fractions were then collected from the size-exclusion column, concentrated to 0.2 to 1 mg/ml using a 30K MWCO concentrator, flash frozen in small aliquots and stored at −80 °C.

#### Dynactin purification

Dynactin was purified as described in (Redwine et al., 2017). HEK293T cells stably expressing p62-Halo-3xFLAG were cultured to ∼95% confluency in 160 × 15 cm plates and then harvested to make pellets. Frozen pellets stored at −80 °C were resuspended in 80 ml of dynactin-lysis buffer (30 mM HEPES (pH 7.4), 50 mM potassium acetate, 2 mM magnesium acetate, 1 mM EGTA, 1 mM DTT, 10% (v/v) glycerol) supplemented with 0.5 mM Mg-ATP, 0.2% Triton X-100, and 1× cOmplete EDTA-free protease inhibitor cocktail tablets (Roche)). Centrifugation (66,000*g*, 30 min, 4 °C) in a Type 70 Ti rotor (Beckman) was performed to clarify the lysate. The supernatant was incubated with 1.5 ml of anti-FLAG M2 affinity gel (Sigma-Aldrich) overnight at 4 °C with rotation. A gravity column was used to collect the beads, which were washed with at least 50 ml of wash buffer (Dynactin-lysis buffer supplemented with 0.1 mM Mg-ATP, 0.5 mM Pefabloc and 0.02% Triton X-100), 100 ml of wash buffer supplemented with 250 mM potassium acetate, and then washed again with 100 ml of wash buffer. Elution was performed by incubating the beads with 1 ml elution buffer (wash buffer with 2 mg/ml of 3xFlag peptide). Dynactin was then diluted to 2 ml in Buffer A (50 mM Tris-HCl (pH 8.0), 2 mM magnesium acetate, 1 mM EGTA, and 1 mM DTT) and loaded onto a MonoQ 5/50 Gl column (Cytiva) at a flow rate of 1 ml/min. Buffer B (50 mM Tris-HCl (pH 8.0), 2 mM magnesium acetate, 1 mM EGTA, 1 mM DTT, 1 M potassium acetate) was run over 26 column volumes as a linear gradient from 35% to 100%. Purified dynactin that eluted in fractions between 75 to 80% Buffer B were collected, pooled, buffer exchanged into GF150 buffer supplemented with 10% glycerol, concentrated to 0.02 to 0.1 mg/ml, flash frozen in small aliquots and stored at −80 °C.

#### Adaptor expression

All parent, chimeric and truncated adaptor constructs containing N-terminal HaloTags were expressed and purified from bacteria, as described (Garrott et al., 2023). All adaptor constructs were transformed into competent BL-21[DE3] cells (New England Biolabs) and were grown in 1.5 L of LB media in a 37 °C shaker until an OD _600_ of 0.4 to 0.6 was reached. Protein expression was then induced with 0.1 mM IPTG and the cultures were grown at 16 °C overnight. The cultures expressing the adaptor were then pelleted by centrifugation, flash-frozen, and stored at −80 °C until needed.

##### Adaptor purification

All adaptor purification steps were carried out at 4 °C, unless otherwise noted. Frozen pellets from 1.5 L of bacterial culture were resuspended in 40 ml of adaptor-lysis buffer (30 mM HEPES pH 7.4, 50 mM potassium acetate, 2 mM magnesium acetate, 1 mM EGTA, 1 mM DTT and 0.5 mM Pefabloc, 10% (v/v) glycerol) supplemented with 1× cOmplete EDTA-free protease inhibitor cocktail tablets (Roche) and 1 mg/ml lysozyme. After resuspension, the slurry was incubated on ice for 30 min before sonication and the lysate was clarified by centrifugation (66,000*g* for 30 min) in Type 70Ti rotor (Beckman). 2 ml of IgG Sepharose 6 Fast Flow beads (Cytiva), equilibrated in adaptor-lysis buffer, were then incubated with the supernatant for 2 h with rotation. The bead-protein mixture was then passed through a gravity column, followed by washing with adaptor-lysis buffer supplemented with 150 mM potassium acetate and 50 ml of adaptor-TEV buffer (50 mM Tris-HCl pH 8.0, 150 mM potassium acetate, 2 mM magnesium acetate, 1 mM EGTA, 1 mM DTT, 0.5 mM Pefabloc and 10% (v/v) glycerol). The washed beads were then incubated overnight in 15 ml of adaptor-TEV buffer supplemented with up to 0.2 mg/ml TEV protease with rotation. Cleaved proteins were collected, concentrated to 1 ml with a 30K MWCO concentrator (Millipore-Sigma), filtered using an Ultrafree-MC VV filter (Millipore-Sigma), diluted to 2 ml in Buffer A (30 mM HEPES pH 7.4, 50 mM potassium acetate, 2 mM magnesium acetate, 1 mM EGTA, 10% (v/v) glycerol and 1 mM DTT) and injected into a 1mL HiTrapQ (Cytiva) or MonoQ 5/50 Gl column (Cytiva) at 1 m/min. Buffer B (30 mM HEPES pH 7.4, 1 M potassium acetate, 2 mM magnesium acetate, 1 mM EGTA, 10% (v/v) glycerol and 1 mM DTT) was run as a linear gradient from 0 to 100% over 26 column volumes and peak fractions containing Halo-tagged adaptors were pooled for concentration. The proteins collected from the ion-exchange column were further diluted to 0.5 ml in GF150 buffer and injected onto a Superdex 200 10/300 column (Cytiva) equilibrated with GF150 buffer. The column was run at a flow rate of 0.5 ml/min before collecting peak fractions. The pooled fractions were then buffer exchanged into GF150 buffer supplemented with 10% glycerol.

#### Adaptor labeling

To label adaptors, concentrated protein fractions were incubated with 15-fold excess of HaloTag-Ligand conjugated TMR or JF646 (Promega) at room temperature for 10 min. A Micro Bio-Spin desalting column preequilibrated with GF150 buffer supplemented with 10% glycerol was used then used to remove excess dye. Proteins were then flash-frozen in small aliquots and stored at −80 °C.

#### Protein quantification

Densitometry analysis of SDS-PAGE gels with a standard curve generated from known quantities of bovine serum albumin was used to calculate the concentrations of dynein, dynactin, Lis1 and all adaptors as described (Gillies et al., 2025). To determine labeling efficiency of each protein, first the absorbance (at 543 nm for TMR labeled protein and 646 nm for JF646 labeled protein) was determined with a DeNovix DS-11 instrument. Then, dye concentrations were calculated using their extinction coefficients: 87,000 M^−1^ cm^−1^ for TMR, and 152,000 M^−1^ cm^−1^ for JF646. Finally, the ratio of molar concentration of dye in each sample to the molar concentration of protein (from densitometry) was used to calculate labeling efficiency.

### Peroxisome relocalization assay

#### Mammalian cell culture

Human osteosarcoma (U2OS) cells were cultured with 1× DMEM (Corning) with 10% FBS (Gibco) and 1% penicillin/streptomycin (Gibco) at 37°C with 5% CO_2_ in a humidified incubator. Cells were passaged in regular intervals (3 – 4 days) using 0.25% trypsin-EDTA (Thermo Fisher Scientific) followed by a 5 min incubation at 37°C to facilitate detachment from the plate. The cells were tested for *Mycoplasma* contamination using the MycoAlert (Lonza) detection kit every 3-6 months.

#### Transfection

#1.5 22 × 22 mm coverslips (Corning) were washed in 100% ethanol and allowed to dry vertically in a 6 -well dish, before coating them with 20 µg/mL PDL diluted in Dulbecco’s phosphate-buffered saline (DPBS; Gibco). The coverslips were then incubated at 37°C for 45 minutes, washed with DPBS and 1× DMEM. 4.5× 10^4^ U2OS cells were plated per well on the PDL-coated coverslips and allowed to adhere overnight. To assess the ability of the adaptors to cluster peroxisomes, cells were then transfected with 500 ng of either StrepII-Halo-NINL^1-702^-V5-FRB (NINL), or StrepII-Halo-NINL^1-275^-BicD2^58-598^-V5-FRB (NB) plasmids using Lipofectamine LTX and PLUS reagents (Thermo Fisher Scientific) in 1X OptiMEM (Gibco) according to manufacturer’s instructions. The cells were allowed to grow for 48h after transfection to facilitate protein expression.

#### Immunofluorescence

*Fixed-cell imaging:* 48h after transfection, the expressed Halo-tagged adaptors were labeled by incubating live cells with 10 nM Janelia Fluor 549 HaloTag Ligand (Promega), diluted in 1× DMEM, for 15-30 minutes at 37°C with 5% CO_2_ in a humidified incubator. Cells were then washed twice with PBS and treated with 1μM A/C Heterodimerizer for 0, 10 or 30 minutes at 37°C with 5% CO_2_ in a humidified incubator. Cells were fixed immediately after treatment with the Heterodimerizer with 4% paraformaldehyde (PFA) for 15 min at room temperature. After fixation, cells were washed with PBS twice and blocked with blocking buffer (0.3% Triton X-100 and 5% normal goat serum (Cell Signaling Technology) in PBS) for 1 h at room temperature in the dark. The cells were washed thrice with PBS before incubation with the primary antibody. Chicken α-GFP (Abcam, RRID: AB_300798), the primary antibody used to stain GFP on peroxisomes, was diluted to 1.25 µg/mL in antibody dilution buffer (0.1% Triton X-100 and 0.5% BSA in 1× PBS), incubated with fixed cells for 1 h in the dark at room temperature, after which the coverslips were washed off thrice with PBS. The secondary antibody, α-chicken-488 (Abcam, RRID: AB_2827653) was diluted to 2 µg/mL in antibody dilution buffer along with 1× F actin probe, phalloidin-647 (Cayman Chemical). The cells were incubated with the mixture of secondary antibody and phalloidin probe and protected from light for 1 h at room temperature. Stained coverslips were then washed thrice with PBS and treated with 1 µg/mL DAPI (Invitrogen) diluted in PBS for 5 min in the dark. The coverslips were then washed four times with PBS and mounted onto glass slides using ProLong Gold Antifade Mountant (Thermo Fisher Scientific). Finally, the slides were cured for 24 h in the dark and sealed with clear nail polish before imaging.

#### Confocal microscopy data acquisition

Cell imaging was performed with an inverted microscope (Nikon, Ti2-E Eclipse) with a 60× 1.40 N.A. oil immersion objective (Nikon, Apo). The microscope was equipped with a LUNF-XL laser launch (Nikon), with 405, 488, 561, and 640nm laser lines. The excitation path was filtered using an appropriate quad band-pass filter cube (Chroma). The emission path was filtered through appropriate emission filters (Chroma) located in a highspeed filter wheel (Finger Lakes Instrumentation). Emitted signals were detected on Photometrics Prime 95B sCMOS Camera. Image acquisition was controlled by NIS-Elements Advanced Research software (Nikon).

#### Peroxisome clustering: fixed-cell imaging and analysis

Fixed cells that had their nucleus, peroxisome, adaptor and actin stained and were imaged with a 60× objective spinning disk confocal microscope. Full volumes of cells were imaged by first identifying the top and bottom of each cell (defined by the peroxisome channel), then acquiring Z-slices at 0.2 µm steps in all four channels. All analysis was performed in FIJI. To begin, all raw Z-stacks were transformed into Z-projections using the “Maximum intensity projection” tool. Then, the actin channel was screened to determine if a single cell could be defined by thresholding. If not, then the brush tool was used to define a clear single-cell area. The actin channel was then subjected to manual thresholding, and the resulting shape was selected with the wand tool to generate a region of interest (ROI) of the cell area. This ROI was then placed on the adaptor channel to measure the mean intensity of the adaptor within the cell as a proxy for expression. The cell area ROI was then placed on the peroxisome channel, and this image was then subjected to automatic Otsu thresholding. The area of all the peroxisome particles (in the threshold) and the area of the cell were saved. The relative peroxisome coverage of each cell was calculated by dividing the area of peroxisomes by the cell area.

#### Peroxisome motility assay: live-cell imaging and analysis

24h after transfection, cells were plated on glass-bottom dishes (Greiner Bio One) pre-coated with PDL. The expressed Halo-tagged adaptors were labeled by incubating live cells with 10 nM Janelia Fluor 549 HaloTag Ligand (Promega), diluted in 1× DMEM, for 15-30 minutes at 37°C with 5% CO_2_ in a humidified incubator. The cells were then washed twice with PBS. Live-cell imaging medium (Gibco) was added to the dishes, and the cells were imaged using a 60× objective spinning disk confocal microscope (Nikon, Ti2-E Eclipse). The intensity of JF549 signal was captured using the 561-laser line and then the peroxisomes were imaged for 2 minutes at 2 frames per second, and 100ms exposure using the 488-laser line. 1uM of A/C Heterodimerizer (Takara Bio) was used to induce the dimerization of FKBP-FRB. 4 minutes after addition of the heterodimerizer, movies capturing peroxisome movement (in cells expressing the adaptor) were collected for up to 12 minutes. All analyses were performed on FIJI. To begin, the mean intensity of the adaptor within the cell was used as a proxy for expression. The adaptor channel was also used to define and measure the cell area. Then, all frames in a movie were transformed into time-projections using the “Hyperstack” tool. This was used to manually trace peroxisome tracks in a cell, all of which were saved in the ROI manager. ImageJ macros (as described (Roberts et al., 2014) were used to generate kymographs of these peroxisome track ROIs from these movies. Velocity (μm/sec) and run length (μm) were also measured for these peroxisomes from the kymographs using custom ImageJ macros described (Roberts et al., 2014). The number of peroxisomes moving processively in each kymograph was also recorded. The values were then summed up to get the total number of peroxisomes moving processively in a cell and then divided by the total cell area.

### Structure predictions

To predict the structures of parent and chimeric adaptors, we used the COSMIC2 web platform to run ColabFold using Colab Notebook (Mirdita et al., 2022; Jumper et al., 2021; Cianfrocco et al., 2017). The structures were visualized using ChimeraX (UCSF) (Pettersen et al., 2021). In the absence of high confidence predicted intramolecular contacts (from PAE plots), the models were manually ‘unfolded’ using ChimeraX (UCSF) and *Coot* (Emsley et al., 2010).

### Single-molecule TIRF microscopy

Single-molecule imaging experiments were all performed on an inverted (Nikon, Ti2-E Eclipse) microscope with a 100×, 1.49 N A. oil immersion objective (Nikon, Apo). The microscope was equipped with a LUNF-XL laser launch (Nikon) containing 405 nm, 488 nm, 561 nm, and 640 nm lasers. An appropriate quad bandpass filter cube (Chroma) was used to filter the excitation path. The emission path was filtered through appropriate emission filters (Chroma) located in a high-speed filter wheel (Finger Lakes Instrumentation). All emission signals were detected on an iXon Ultra 897 electron-multiplying CCD camera (Andor Technology). NIS Elements Advanced Research software (Nikon) was used to control the microscope and image acquisition.

### Motility assay and analysis

#### Data collection

Single-molecule motility experiments were performed in flow chambers assembled as described previously (DeSantis et al., 2017) using No. 1-1/2 coverslips (Corning) functionalized with PEG-biotin. 20 μM bovine tubulin (with ∼10% biotin-tubulin and ∼10% Alexa-488 tubulin) was polymerized into microtubules at 37 °C for 30 min before being stabilized with 20 μM taxol. To assemble the flow chambers, 1 mg/ml streptavidin in assay buffer (30 mM HEPES [pH 7.4], 2 mM magnesium acetate, 1 mM EGTA, 10% glycerol, 1 mM DTT, and 20 μM taxol) was first flowed in and incubated for 3 min before washing the chambers twice with 20 μl of assay buffer. Next, a fresh-dilution of taxol-stabilized microtubules (final concentrations ∼0.1–0.4 μM) was flowed in and incubated for 3 min. The chambers were again washed twice with assay buffer supplemented with 1 mg/ml casein. To assemble dynein-dynactin-adaptor complexes, dynein (Final concentration 0.5-0.75 nM), dynactin, and the adaptor were mixed at a 1:2:10 M ratio and incubated for 10 min on ice. These complexes were incubated for another 10 min on ice with Lis1(final concentration 50 nM) (or GF150- to buffer match) and assay buffer supplemented with 1 mg/ml casein. Immediately before imaging, the samples were mixed with assay buffer supplemented with 1 mg/ml casein, 71.5 mM β-mercaptoethanol, 0.05 mg/ml glucose catalase, 1.2 mg/ml glucose oxidase, 0.4% glucose, and 2.5 mM Mg-ATP. Samples were then introduced into the flow chamber and imaged immediately. Microtubules were imaged first by taking a snapshot of a single field of view. Dynein (labeled with JF646 or TMR) was imaged every 300 msec for 3 min. The microtubule channel was imaged again at the end of the image acquisition to assess stage drift. Movies showing drift were excluded.

#### Analysis

ImageJ macros were used to generate kymographs from movies as described (Roberts et al., 2014). Dynein’s landing rate was determined per microtubule according to the following equation: landing rate=Νpe/(lm·[dynein]·t), where Νpe is the number of processive events, lm is the microtubule length (in μm) and t is the length of the movie (in minutes). Velocity (μm/sec) and run length (μm) were also measured for these processive events from the kymographs using custom ImageJ macros described (Roberts et al., 2014). The BicD2 data shown in Figure 2E and Figure S2E-F was analyzed using Kymobutler. Runs shorter than eight pixels and bright protein aggregates (defined as molecules 4 times brighter than the background) were all excluded from the analysis. Relative motile propensity was calculated by taking all the median values for either velocity, run length and landing rate, subtracting the minimum value, and then dividing by the maximum value to give values ranging from 0-1. These were then summed to give RMP values ranging from 0-3. Statistical analyses performed within data sets were performed in Prism10 (GraphPad). Correlation analysis was performed in Matlab (Mathworks).

#### Reproducibility

To ensure reproducibility most data was collected with at least two preps of dynein and two preps of adaptor. The only exceptions to this are the BicD2 truncation experiments presented in Figure 2 and the two-color dynein experiment and the labelled adaptor experiment presented in Figure 5, for which only one prep of each was used. We have observed that if the components are left on ice for too long the landing rate of dynein can decrease. To combat this, we limited the number of samples collected at one time, and we changed the order that samples are collected within a given experiment. The density of microtubules attached to the coverslip can also influence landing rate, and so chambers were visually confirmed to have a similar microtubule density prior to use. At least one rep of each chimera’s motility was collected alongside the parent adaptors to confirm comparable activity. The peroxisome relocalization data in U2OS cells, presented in Figure 1 and Figure 5, were independently repeated three times to ensure reproducibility.

### Equilibrium binding experiments

To measure the binding of dynein to adaptor in solution, adaptors were coupled to 15 μl of Magnetic Halotag beads (Promega) in 2 ml Protein Lo Bind Tubes (Eppendorf) following the method outlined below. Beads were first washed twice with 1 ml of GF150 supplemented with 10% glycerol and 0.1% NP40. The adaptors were diluted in this buffer from 0-600nM. 25 μl of diluted adaptors were added to the beads and shaken gently for 1 h. 20 μl of supernatant was then run on an SDS-PAGE gel to confirm complete depletion of the protein by the beads. The protein-conjugated beads were washed once with 1 ml GF150 with 10% glycerol and 0.1% NP40, followed by one more wash with 1 ml of binding buffer (28.75 mM HEPES [pH 7.4], 1.5 mM magnesium acetate, 0.75 mM EGTA, 0.25 mM MgCl_2_, 37.5 mM KCl, 10% glycerol, 1 mM DTT, 1 mg/ml casein, 0.1% NP40, 1 mM ADP). 5 nM dynein was diluted such that the final buffer composition was the same as the binding buffer. 25 μl of the diluted dynein was then added to the beads and gently shaken for 45 min. Finally, 20 μl of the supernatant was removed, and mixed with 6.67 μl of NuPAGE LDS Sample Buffer (4×) and 1.33 μl of Beta-mercaptoethanol. The samples were boiled for 10 min before running on a 4-12% NuPAGE Bis-Tris gel. Depletion was determined using densitometry in ImageJ. To measure dynein-dynactin-adaptor complex formation the same protocol was used with the following changes: 35 μl of NEBExpress Ni-NTA Magnetic Beads (NEB) were conjugated to 90nM of adaptor, and 5nM dynein and 5nM dynactin was in solution during the binding step.

### Statistical analysis

GraphPad Prism was used to perform all statistical analyses. The exact statistical tests used, *n* value, and number of replicates are noted in the figure legend. Statistical significance: ns, p > 0.05; *, p ≤ 0.05; **, p ≤ 0.01; ***, p ≤ 0.001; ****, p ≤ 0.0001. For correlations the p-value cutoff was corrected for the ten comparisons made, so the cutoff are ns, p > 0.005; *, p ≤ 0.005; **, p ≤ 0.001; ***, p ≤ 0.0001.

## Supplementary figures

**Figure S1:**
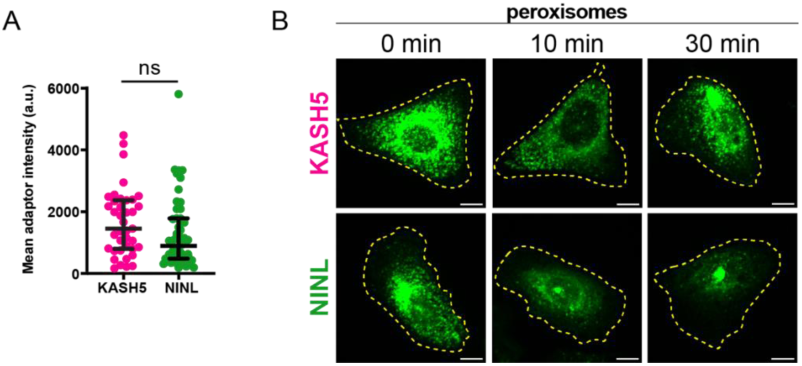
**A.** Plot showing the mean intensity of adaptors in cells at 0 min. Error bars indicate median ± interquartile range. Statistical analysis was performed using a Mann-Whitney test. n = 39 cells for KASH5 and 50 cells for NINL. p-value can be found in Supplemental file 1. **B.** Fluorescence microscopy images of fixed U2OS cells expressing KASH5 (top panel) or NINL (bottom panel) and stained for peroxisomes with anti-GFP at 0,10 and 30 min after treatment with A/C Heterodimerizer. Yellow dashed line marks the boundary of the cell. Scale bar = 10µm.

**Figure S2:**
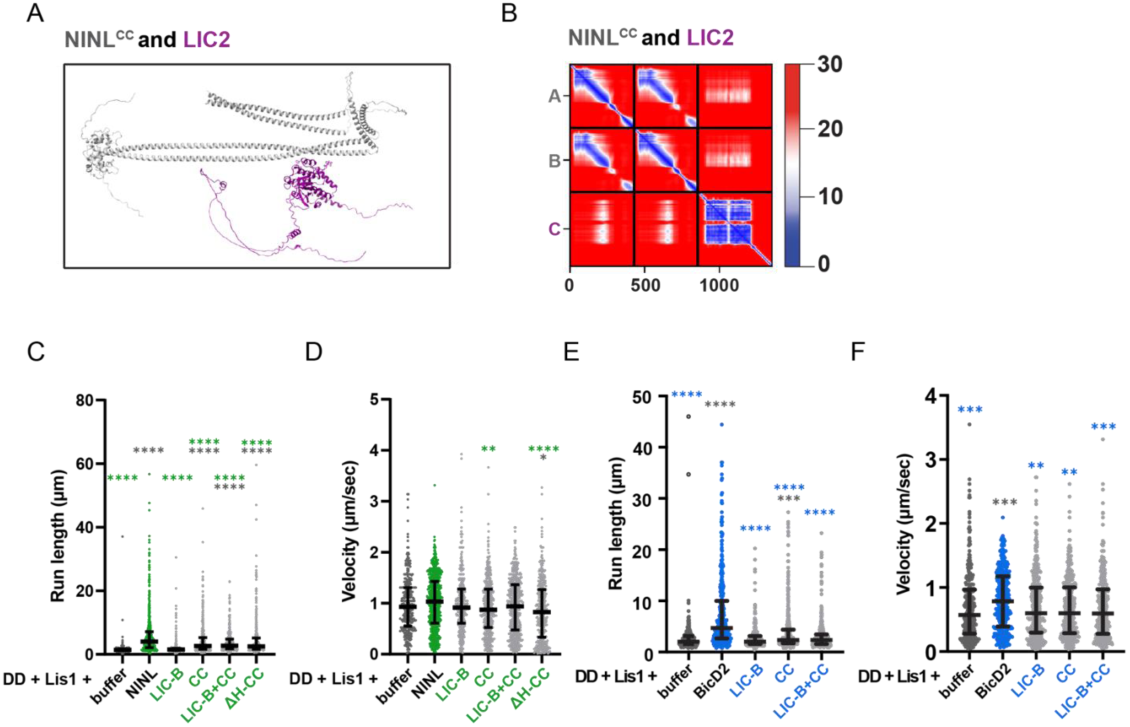
**A.** ColabFold model showing NINL^CC^ (gray) with dynein LIC2 (purple). **B.** Predicted aligned error (PAE) plot of the model in A, where blue indicates high predictive confidence and red indicates low predictive confidence. **C.** Single-molecule run lengths (µm) of DD-Lis1 complexes incubated with either buffer (dark gray dots), NINL (green dots), NINL^LIC-B^, NINL^CC^, a combination of NINL^LIC-B^ and NINL^CC^, or NINL^ΔH-CC^. Error bars are median ± interquartile range. Statistical analysis was performed using a Kruskal-Wallis with Dunn’s multiple comparisons test. Samples are compared to NINL (green stars) or buffer (dark gray stars). p-values and n can be found in Supplemental file 1. **D.** Single-molecule velocity (µm/sec) of DD-Lis1 complexes incubated with either buffer (dark gray dots), NINL (green dots), NINL^LIC-B^, NINL^CC^, a combination of NINL^LIC-B^ and NINL^CC^, or NINL^ΔH-CC^. Error bars are median ± interquartile range. Statistical analysis was performed using a Kruskal-Wallis with Dunn’s multiple comparisons test. Samples are compared to NINL (green stars) or buffer (dark gray stars). p-values and n can be found in Supplemental file 1. **E.** Single-molecule run lengths (µm) of DD-Lis1 complexes incubated with either buffer (dark gray dots), BicD2 (blue dots), BicD2^LIC-B^, BicD2^CC^, or a combination of BicD2^LIC-B^ and BicD2^CC^. Error bars are median ± interquartile range. Statistical analysis was performed using a Kruskal-Wallis with Dunn’s multiple comparisons test. Samples are compared to BicD2 (blue stars) or buffer (dark gray stars). p-values and n can be found in Supplemental file 1. **F.** Single-molecule velocity (µm/sec) of DD-Lis1 complexes incubated with either buffer (dark gray dots), BicD2 (blue dots), BicD2^LIC-B^, BicD2^CC^, or a combination of BicD2^LIC-B^ and BicD2^CC^. Error bars are median ± interquartile range. Statistical analysis was performed using a Kruskal-Wallis with Dunn’s multiple comparisons test. Samples are compared to BicD2 (blue stars) or buffer (dark gray stars). p-values and n can be found in Supplemental file 1.

**Figure S3.1:**
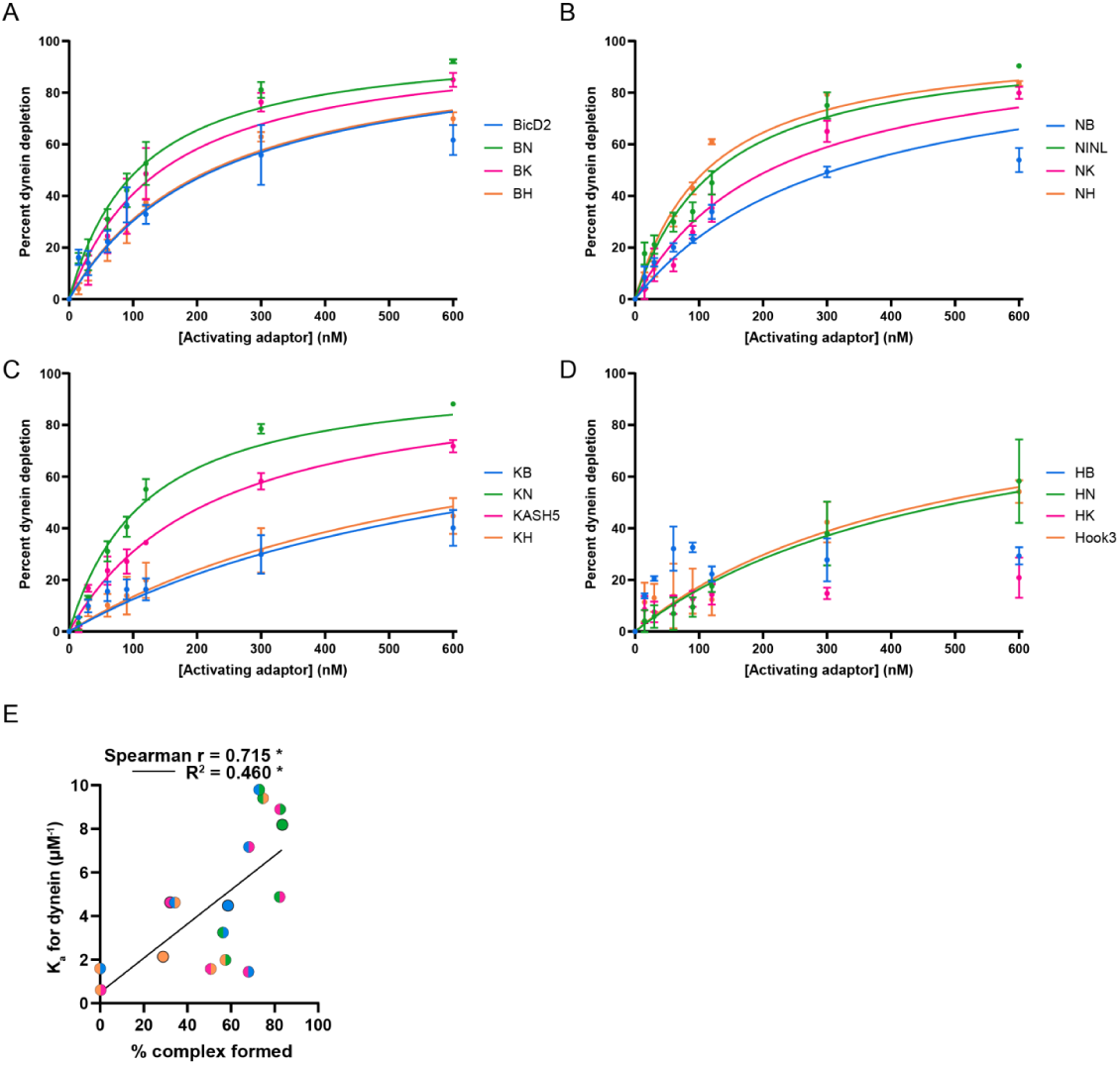
**A-D.** Plots showing the binding curve between adaptors (grouped by the identity of the LIC-B domain) and dynein. The LIC-B domain originates from **(A)** BicD2, **(B)** NINL, **(C)** KASH5 and **(D)** Hook3. Error bars are mean ± SEM n = 3 for all samples. In each plot, the color of the binding curve indicates the parent adaptor from which the coiled-coil originates. **E.** Scatter plot showing the K_a_ (µM^-1^) of different adaptors to dynein and the percentage of dynein-dynactin-adaptor complexes formed with the respective adaptors. Each circle represents an adaptor, where the color in the first half of the circle represents the identity of the LIC-B domain and the color in the second half of the circle represents the identity of the coiled-coil domain. Spearman’s Correlation coefficient (r) and Pearson’s coefficient of determination (R^2^) are indicated. p values can be found in Supplemental file 1.

**Figure S3.2:**
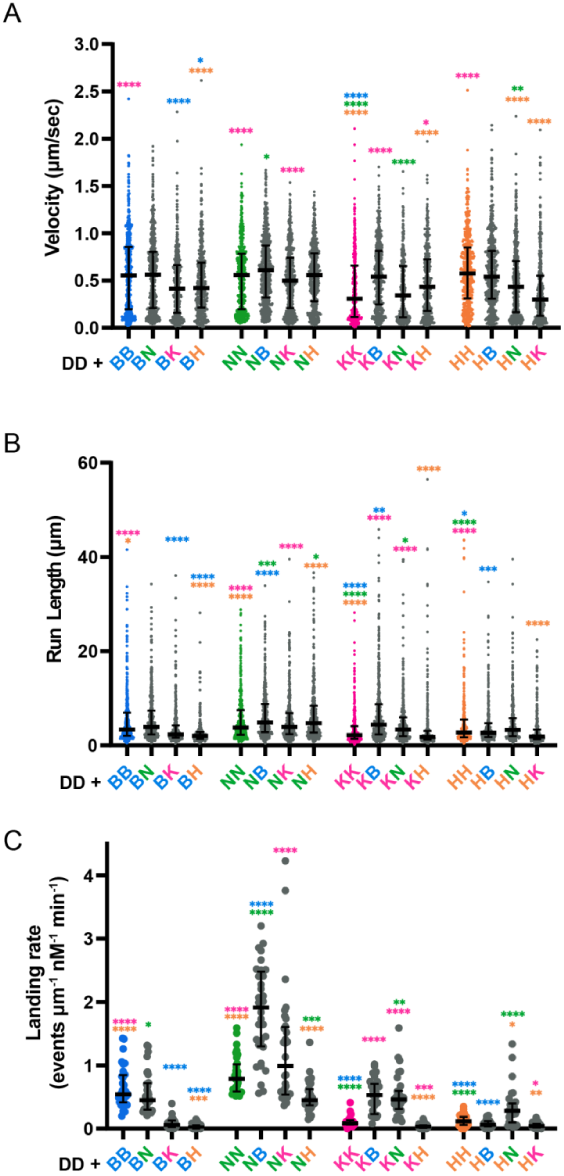
**A.** Single-molecule velocity (µm/sec) of activated dynein-dynactin-adaptor complexes. Error bars are median ± interquartile range. All chimeric adaptors are shown in gray, and parent adaptors are shown in their respective colors (BicD2- blue, NINL- green, KASH5- pink, Hook3- orange). The color of the asterisk indicates the parent adaptor from which it is significantly different. Statistical analysis was performed using a Kruskal-Wallis with Dunn’s multiple comparisons test. p-values and n can be found in Supplemental file 1. **B.** Single-molecule run lengths (µm) of activated dynein-dynactin-adaptor complexes. Error bars are median ± interquartile range. All chimeric adaptors are shown in gray, and parent adaptors are shown in their respective colors (BicD2- blue, NINL- green, KASH5- pink, Hook3- orange). The color of the asterisk indicates the parent adaptor from which it is significantly different. Statistical analysis was performed using a Kruskal-Wallis with Dunn’s multiple comparisons test. p-values and n can be found in Supplemental file 1. **C.** Landing rates of processive complexes reported as events per micrometer of microtubule per nanomolar dynein per minute. Error bars are median ± interquartile range. All chimeric adaptors are shown in gray, and parent adaptors are shown in their respective colors (BicD2- blue, NINL- green, KASH5- pink, Hook3- orange). The color of the asterisk indicates the parent adaptor from which it is significantly different. Statistical analysis was performed using a Brown-Forsythe and Welch ANOVA with Dunnett’s T3 multiple comparisons test. p-values and n can be found in Supplemental file 1.

**Figure S3.3:**
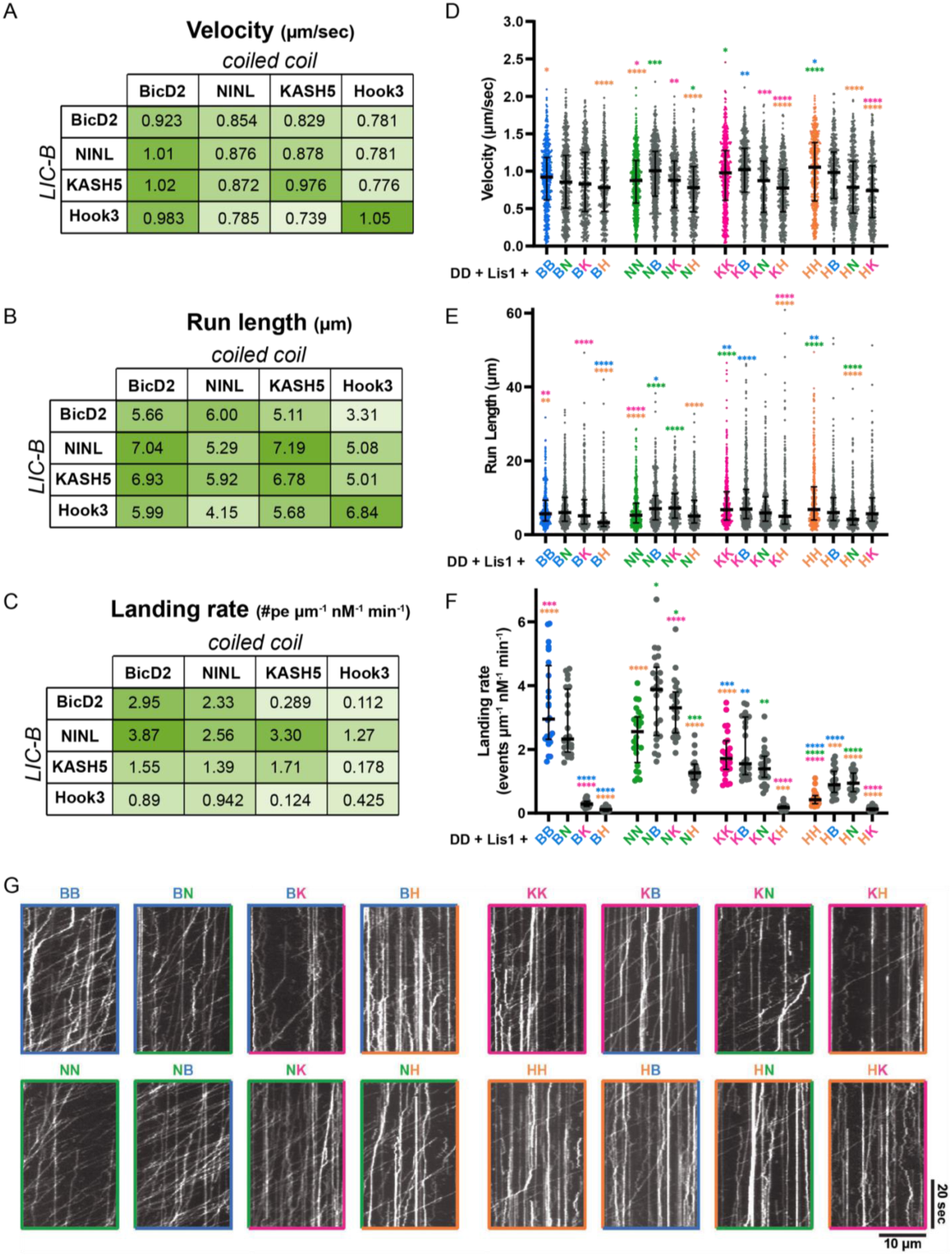
**A-C.** Median single-molecule **(A)** velocity (µm/sec), **(B)** run length (µm), and **(C)** landing rate (processive events/ µm nM min) of dynein complexes formed with different parent and chimeric adaptors in the presence of Lis1 are shown as tables, where each cell represents an adaptor. The identity of the LIC-B domain in an adaptor is indicated by the columns on the left, and the identity of the coiled-coil is indicated by the top rows. The range of values (velocity, run length or processive landing rate) observed with different adaptors is reflected by the color gradient in each table, where lighter green shades represent lower median values and darker green shades indicate higher median values. **D.** Single-molecule velocity (µm/sec) of activated dynein-dynactin-adaptor complexes formed in the presence of Lis1. Error bars are median ± interquartile range. All chimeric adaptors are shown in gray, and parent adaptors are shown in their respective colors (BicD2- blue, NINL- green, KASH5- pink, Hook3- orange). The color of the asterisk indicates the parent adaptor from which it is significantly different. Statistical analysis was performed using a Kruskal-Wallis with Dunn’s multiple comparisons test. p-values and n can be found in Supplemental file 1. **E.** Single-molecule run lengths (µm) of activated dynein-dynactin-adaptor complexes formed in the presence of Lis1. Error bars are median ± interquartile range. All chimeric adaptors are shown in gray, and parent adaptors are shown in their respective colors (BicD2- blue, NINL- green, KASH5- pink, Hook3- orange). The color of the asterisk indicates the parent adaptor from which it is significantly different. Statistical analysis was performed using a Kruskal-Wallis with Dunn’s multiple comparisons test. p-values and n can be found in Supplemental file 1. **F.** Landing rates of processive complexes formed in the presence of Lis1 reported as events per micrometer of microtubule per nanomolar dynein per minute. Error bars are median ± interquartile range. All chimeric adaptors are shown in gray, and parent adaptors are shown in their respective colors (BicD2- blue, NINL- green, KASH5- pink, Hook3- orange). The color of the asterisk indicates the parent adaptor from which it is significantly different. Statistical analysis was performed using a Brown-Forsythe and Welch ANOVA with Dunnett’s T3 multiple comparisons test. p-values and n can be found in Supplemental file 1. **G.** Example kymographs of dynein complexes formed with parent and chimeric adaptors in the presence of Lis1.

**Figure S4:**
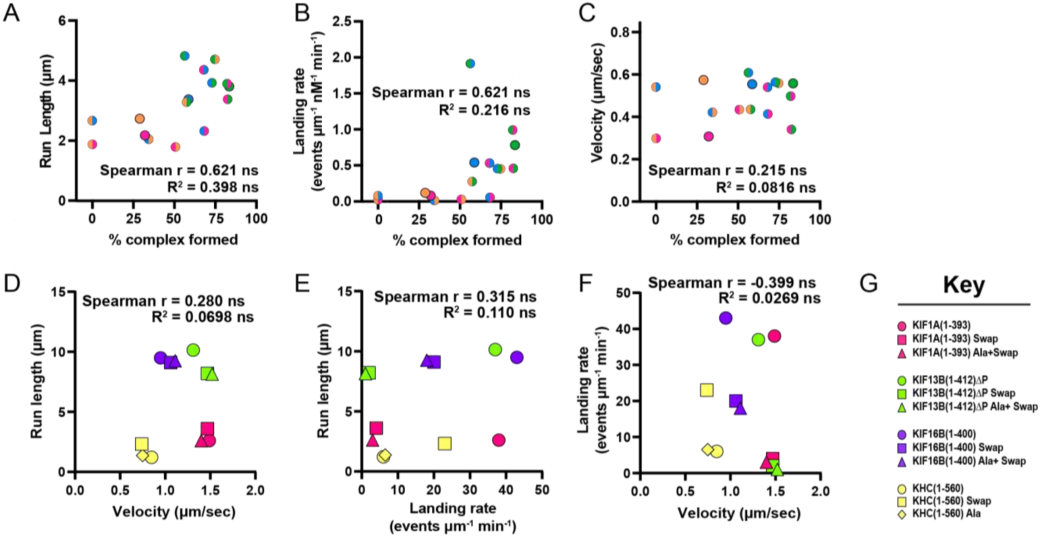
**A-C.** Scatter plot showing percentage of dynein-dynactin-adaptor complexes formed (x-axes) with different adaptors and their corresponding **(A)** median run length (µm) **(B)** median landing rate (processive events/µm nM min) and **(C)** median velocity (µm/sec). Spearman’s Correlation coefficient (r) and Pearson’s coefficient of determination (R^2^) are indicated. p-values can be found in Supplemental file 1. **D-F**. Relation between **(D)** run length (µm) and velocity (µm/sec), **(E)** run length (µm) and landing rate (events/µm min), and **(F)** landing rate (events/µm min) and velocity (µm/sec) of different kinesin motors and their mutants as measured in (Soppina and Verhey, 2014b). Correlation coefficient (r) and Pearson’s coefficient of determination (R^2^) are indicated. p-values can be found in Supplemental file 1. **(G)** Key indicating the kinesin protein and mutation in D-F.

**Figure S5:**
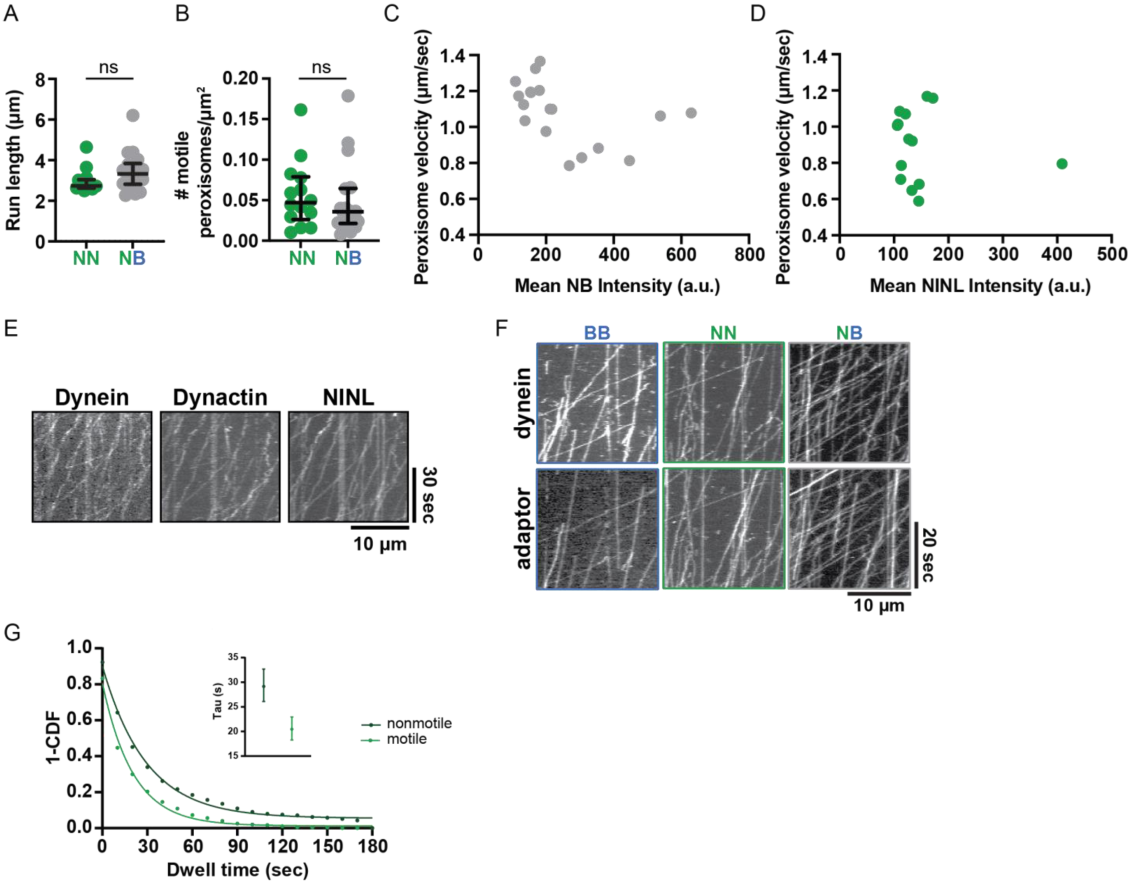
**A.** Run length (µm) of processively moving peroxisomes in U2OS cells expressing NN (green) or NB (gray). Error bars are median ± interquartile range. Statistical analysis was performed using a Mann-Whitney test. n= 14 cells for NINL and 18 cells for NB. p values can be found in Supplemental file 1. **B.** Number of processively moving peroxisomes per unit cell area in U2OS cells expressing NN (green) or NB (gray). Error bars are median ± interquartile range in A and B. Statistical analysis for A and B was performed using a Mann-Whitney test. n= 14 cells for NN and 18 cells for NB. p values can be found in Supplemental file 1. **C-D.** Scatter plot showing the median velocity (µm/sec) of moving peroxisomes (x-axes) and the adaptor intensity (y-axes) in cells expressing (C) NB or (D) NINL. **E.** Kymographs showing the colocalization between dynein-488, dynactin-647 and NINL-TMR. **F.** Example kymographs of dynein-dynactin-adaptor complexes formed with BB, NN or NB, in which both dynein (top panel) and adaptor (bottom panel) are labelled. **G.** 1- cumulative density function for the dwell times of dynein-dynactin-NINL complexes presented in C that are motile (light green) or nonmotile (dark green). Tau values from the exponential decay are shown in the inset.

**Figure S6:**
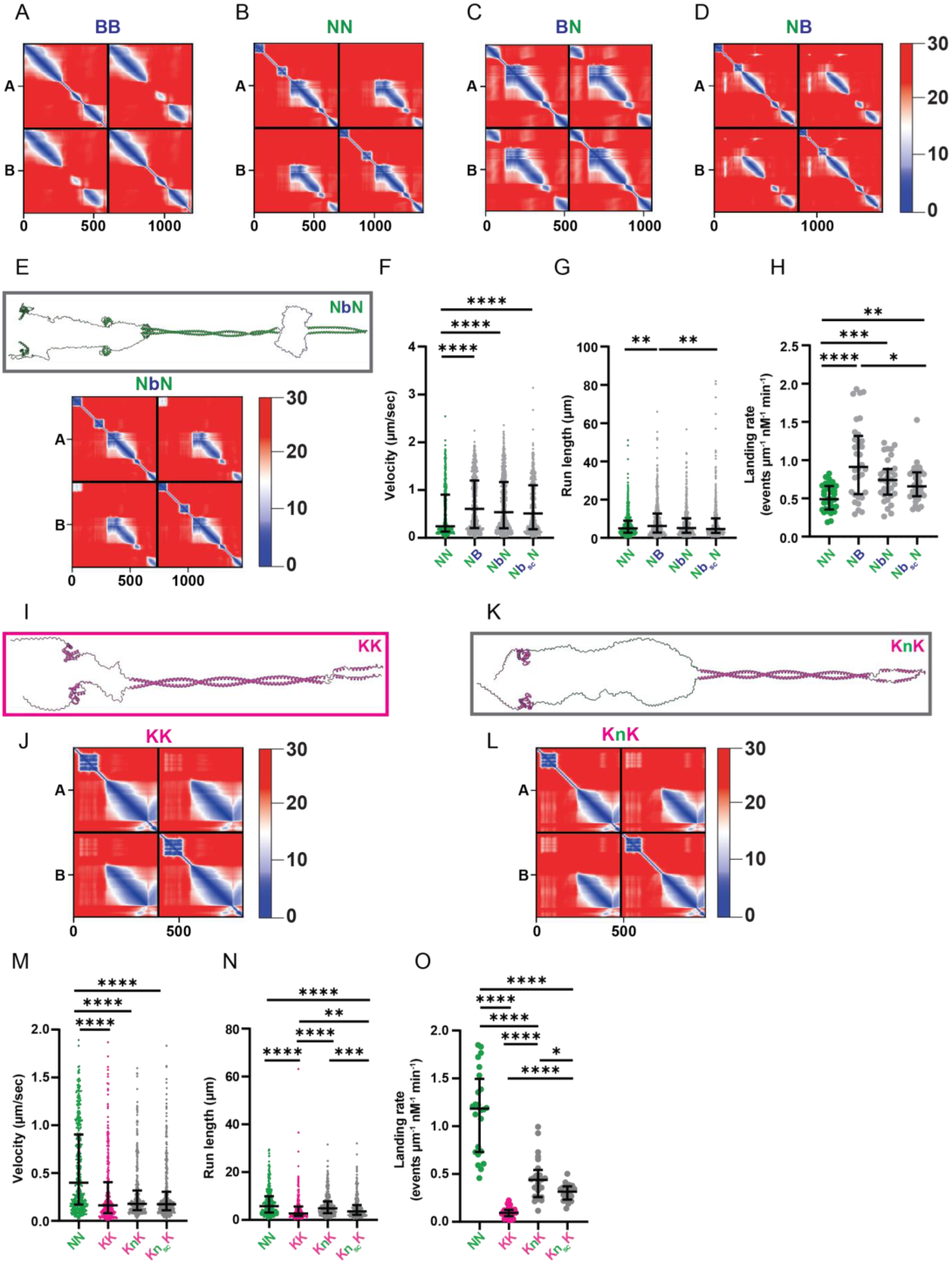
**A-D.** Predicted aligned error (PAE) plots for the corresponding ColabFold predictions of **(A)** BicD2, **(B)** NINL, **(C)** BN, and **(D)** NB, where blue indicates high predictive confidence and red indicates low predictive confidence. **E.** ColabFold prediction of NbN and its corresponding PAE plot. Flexible regions (with low reliability) in these structures were artificially ‘linearized’ using *Coot* for better visualization. **F-H. (F)** Velocity (µm/sec), and **(G)** run length (µm) and **(H)** landing rate (processive events per micrometer of microtubule per nanomolar of dynein per minute), of dynein complexes formed with NN (green), NB, NbN and Nb_sc_N. Error bars indicate median ± interquartile range. Statistical analysis was performed using a Kruskal-Wallis with Dunn’s multiple comparisons test in F and G, and a Brown-Forsythe and Welch ANOVA with Dunnett’s T3 multiple comparisons test in H. p-values and n can be found in Supplemental file 1. **I-L.** ColabFold prediction of **(I)** KASH5 with **(J)** its corresponding PAE plot and **(K)** KnK with **(L)** its corresponding PAE plot. Flexible regions (with low reliability) in these structures were artificially ‘linearized’ using *Coot* for better visualization. **M-O. (M)** Velocity (µm/sec), **(N)** run length (µm) and **(O)** landing rate (processive events per micrometer of microtubule per nanomolar of dynein per minute), of dynein complexes formed with NN (green), KK (pink), KnK and Kn_sc_K. Error bars indicate median ± interquartile range. Statistical analysis was performed using a Kruskal-Wallis with Dunn’s multiple comparisons test in M and N, and a Brown-Forsythe and Welch ANOVA with Dunnett’s T3 multiple comparisons test in O. p-values and n can be found in Supplemental file 1.

